# The importance of postzygotic barriers at the early stages of speciation in trees

**DOI:** 10.64898/2026.02.23.707326

**Authors:** Elizabeth A. Stacy, Alicia M. Rhoades, Kevin W. Brinck, April H. Wallace

**Author notes:** Author for correspondence: Elizabeth A. Stacy.

## Abstract

Recent reviews of isolating barriers in plants conclude that prezygotic barriers play an outsized role in plant speciation; yet these conclusions derive overwhelmingly from studies of sympatric, perennial herbs in temperate zones, and at later stages of speciation. Trees possess several traits that are expected to influence barrier evolution, including prolonged generation times and reproduction, predominant outcrossing, and long-distance gene flow. We examined early-evolving post-pollination barriers between ecologically diverged, vegetatively distinct varieties of the tree species, *Metrosideros polymorpha*, that have a common floral morphology and highly overlapping flowering times. We performed controlled crosses between each of Hawaii Island’s four varieties and maternal trees of the high-elevation variety and examined pollen-tube growth, fruit set, seed germination, and seedling phenotype. We then monitored survivorship, maturation rate, and fertility of F_1_ hybrids over ≥8 years alongside parental controls and a fourth F_1_ genotype derived from companion studies. The four F_1_ crosses showed four contrasting patterns and strengths of predominantly postzygotic isolation, including high F_1_ mortality that manifested over several years. Results from this and other tree studies suggest that ecological speciation in trees follows the classical speciation model of early postzygotic barrier formation followed by reinforcement, whenever stable environments promote recurring hybridization.

## Introduction

How isolating barriers form during plant speciation is a long-standing question in evolutionary biology. Reviews of isolating barriers in plants suggest that they likely accumulate differently in different groups (Moyle *et al*., 2004; Rieseberg & Willis, 2007; Christie *et al*., 2022) and that prezygotic barriers likely play an outsized role at the early stages of speciation (Lowry *et al*., 2008), but see Widmer *et al*. (2009). Among prezygotic barriers examined, those associated with extrinsic or ecological factors appear to be most important, including floral isolation, ecogeographic isolation, and immigrant inviability (Rieseberg & Willis, 2007; Christie *et al*., 2022). Meanwhile, the most commonly measured postzygotic barrier, F_1_ viability, varied among taxon pairs and was found to be ineffective overall in halting gene flow (Christie *et al*., 2022). As noted by the authors of these reviews, however, these conclusions derive from a body of work that does not represent the full spectrum of plant life histories, geographies, or stages of speciation. Rather, the majority of taxa comprise sympatric, perennial herbs in temperate zones and at later stages of speciation (Christie *et al*., 2022). When studying isolation above the species level, where multiple barriers are often present (Lowry *et al*., 2008; Widmer *et al*., 2009; Scopece *et al*., 2013), discerning which barriers arose before versus after speciation is difficult. Further, complete prezygotic isolation, if present, precludes the detection of postzygotic mechanisms (Coyne & Orr, 2004). As a result, and because of the greater challenges often involved in characterizing postzygotic barriers, their importance early in speciation may be under-estimated (Coughlan & Matute, 2020). Studies of isolating barriers across a broader range of plant life histories and geographic regions, and focusing on earlier stages of divergence, are needed for a fuller understanding of the barriers that form early, and thus drive, speciation in plants (Lowry, 2012; Ostevik *et al*., 2016).

An important life history that is conspicuously under-represented in studies of isolating barriers in plants is the long-lived perennial, or tree (Baack *et al*., 2015). There are more than 70,000 tree species globally (Cazzolla Gatti *et al*., 2022), yet how these species arise is a conundrum, given the many characteristics of trees that are expected to impede speciation. First, long generation times slow the speciation rate (Petit & Hampe, 2006). Second, because of the high genetic load typical of trees, the vast majority of species are nearly obligately outcrossed (Petit & Hampe, 2006), and third, given their tall stature and high fecundity, trees have a tremendous potential for long-distance gene flow by pollen (e.g., Dick *et al*., 2008; Slavov *et al*., 2009; Kamm *et al*., 2009). As a result, differential local adaptation to contrasting regeneration niches (Poorter, 2007) is unlikely to isolate parapatric tree populations (i.e., through ecogeographic isolation) as effectively as it would for herbaceous plants. Moreover, because of the predominant outcrossing and potential for long-distance dispersal typical of trees (Loveless & Hamrick, 1984; Bawa, 1992), even geographically widespread populations are less often allopatric, as defined by a lack of gene flow. High gene flow across broad spatial scales is evident in the weak genetic structure observed over vast areas for many temperate tree species (e.g., Bodénès *et al*., 1997; Parchman *et al*., 2011), and the resulting large effective population sizes are expected to slow speciation rates (Shang *et al*., 2020). Owing to their long life spans and tremendous, multi-year fecundity, trees can also tolerate periods of reproductive failure - such as the production of ecologically misfit hybrid offspring - without negative consequences for demography (Calvo & Horvitz, 1990; Ashman *et al*., 2004; Petit & Hampe, 2006). As such, the consequences of poor mate choice are less dire for trees than for short-lived plants, thus resulting in weaker selection for isolating barriers. This prediction of weaker isolating barriers among closely related tree species is supported by the greater frequency of natural hybrids in outcrossed, perennial plants compared to other plants in five floras (Ellstrand *et al*., 1996).

Finally, in contrast to the primary role of isolation by floral characters inferred for shorter-lived plants, diversification within most species-rich tree groups appears to be predominantly through vegetative characters, suggesting a secondary role for pollinators in isolation. Canopy trees in northern temperate zones are wind-pollinated, while the floral morphology of animal-pollinated canopy trees elsewhere appears to be largely uniform within the most species-rich groups. These flowers typically attract a community of pollinators rather than a narrow group of specialists [e.g., *Inga*-hummingbirds, hawkmoths, bats (Amorim *et al*., 2013), Southeast Asian Dipterocarps-bees, beetles, thrips (Momose *et al*., 1998); a mix of invertebrate or both invertebrate and vertebrate pollinators in the three most species-rich genera of Myrtaceae: *Syzygium* (Hopper, 1980; Boulter *et al*., 2005; Badou *et al*., 2020), *Eugenia* (Rajkumar *et al*., 2015; Diniz & Buschini, 2015), and *Eucalyptus* (Hingston *et al*., 2004)]. As a result, diversification through selection by specialist pollinators appears to be less likely in trees compared to herbaceous plants, although closer examination of pollinator assemblages may reveal biases of individual species toward individual tree species (Misiewicz *et al*., 2020). An obvious exception to this pattern is the species-specific brood-pollination mutualisms of the hyper-diverse genus *Ficus* (Cruaud *et al*., 2012), where the linkage between plant and pollinator reproduction, selection for strong host discrimination by pollinators, and the short generation time of pollinators relative to the host plant may be sufficient to effect prezygotic isolation among sympatric fig species (Moe *et al*., 2012; Moe & Weiblen, 2012).

Insights into isolating barriers among closely related tree species are limited due to the difficulty of experimenting with very tall, long-lived organisms; however, studies to date have identified a range of pre- and post-zygotic barriers contributing to the maintenance of congeneric species in sympatry. In eucalypts, which may be the most understood trees with respect to isolating barriers, a broad spectrum of mechanisms has been recorded, from diverged flowering times (Orel *et al*., 2025) to pollen-stigma interactions (Dickinson *et al*., 2012), insufficient or aborted pollen tube growth (Ellis *et al*., 1991) sometimes associated with contrasting flower sizes (Gore *et al*., 1990; Larcombe *et al*., 2016a), reduced F_1_ seed germination and/or early seedling survivorship, and abnormal F_1_ seedling phenotypes (Potts *et al*., 2000; Potts & Dungey, 2004).

Early-acting barriers identified in other trees include flowering time, pollen-stigma/style interactions and early postzygotic barriers (Protium subserratum, Misiewicz *et al*., 2020), insufficient or aborted pollen tube growth (oaks, Abadie *et al*., 2012), conspecific pollen precedence (European oaks, Lepais *et al*., 2013), and early embryo failure (American oaks, Williams *et al*., 2001). Where studies have monitored hybrid fitness over several years, however, significant later-acting, intrinsic barriers have also been observed, including reduced growth rates over five years of 17 *Eucalyptus* F_1_ genotypes (Larcombe *et al*., 2016a) and reduced survivorship between two and 10 years after seed germination in three *Eucalyptus* F_1_ genotypes (Lopez *et al*., 2000; Barbour *et al*., 2006; Costa e Silva *et al*., 2012; Larcombe *et al*., 2014).

Further, where second- or later-generation hybrids have been examined, significant reductions in fitness, again often appearing beyond the seedling stage, have been reported, including reduced survivorship and/or fertility (Eucalyptus, Potts & Dungey, 2004; Costa e Silva *et al*., 2012; Larcombe *et al*., 2016b; Metrosideros, Stacy *et al*., 2017). In *Populus* in particular, early-acting barriers appear insufficient to prevent cross-fertilization among species in hybrid zones (Lindtke *et al*., 2012; Roe *et al*., 2014, and references within), and late-acting selection on recombinant hybrids likely maintains species boundaries (Lindtke *et al*., 2014). The long generation times and delayed maturity of trees extend the period over which hybrid incompatibilities are expressed relative to short-lived plants (Levin, 2012; Lindtke *et al*., 2014; Larcombe *et al*., 2016b), and understanding isolation through hybrid weakness or inviability beyond the seedling stage requires long-term study.

Hawaiian *Metrosideros* Banks ex Gaertn (Myrtaceae) is a rare example of an ongoing, predominantly sympatric, adaptive radiation of trees that allows examination of isolating barriers at multiple early stages of speciation (Stacy *et al*., 2014, 2017; Izuno *et al*., 2022). This 3-4- MYO radiation (Wright *et al*., 2000; Percy *et al*., 2008; Dupuis *et al*., 2019; Choi *et al*., 2021) comprises >20 vegetatively distinct races, varieties, and species of woody taxa that collectively dominate native forests of the Hawaiian Islands (Stacy & Sakishima, 2019). *Metrosideros* taxa range from prostrate plants to 30-m trees and sport a broad range of leaf sizes and morphologies, from glabrous to hirsute (Dawson & Stemmermann, 1990). Within continuous *Metrosideros* stands on each island, individual taxa are nonrandomly distributed across environments (Stemmermann, 1983; Dawson & Stemmermann, 1990; Stacy *et al*., 2020), and studies from two islands support a model of phenotypic diversification through differential local adaptation (Morrison & Stacy, 2014; Sur *et al*., 2018; Ekar *et al*., 2019; Stacy & Ostertag, 2025; Merondun *et al*., in prep).

*Metrosideros* on young (<700 KYA; Clague, 1996), volcanically active Hawaii Island comprises just four varieties of the archipelago-wide species, *M. polymorpha*, that together dominate the largest continuous forest on any Hawaiian Island. Within this continuous *Metrosideros* community, the four varieties (or ecotypes; Turesson, 1922; Lowry, 2012) are distributed across four environments (Fig. 1) and hybridize where they co-occur in intermediate environments. Early-successional, pubescent *M. polymorpha* var. *incana* (hereafter incana) occurs from sea level to middle elevations on new lava flows and is replaced over time by the late-successional, glabrous variety, *M. polymorpha* var. *glaberrima* (glaberrima), which dominates older substrates across a broader elevation range (Stemmermann, 1983; Drake & Mueller-Dombois, 1993); pubescent *M. polymorpha* var. *polymorpha* (polymorpha) dominates both early- and late-successional communities at high elevations, and typically-glabrous riparian *M. polymorpha* var. *newellii* (newellii) is endemic to waterways on the windward side of the island (Dawson & Stemmermann, 1990). Whereas incana, glaberrima, and polymorpha have broad, overlapping distributions and likely colonized Hawaii Island from older Maui Nui following the progression rule (Percy *et al*., 2008; Choi *et al*., 2020; Izuno *et al*., 2022), newellii is very narrowly distributed and appears to have emerged in situ from glaberrima through disruptive selection with primary gene flow along a sharp forest-riparian ecotone (Choi *et al*., 2020). These four varieties are distinguished through vegetative characters and are weakly to moderately differentiated at neutral genetic markers (Figs. 1 and 2). All taxa are highly dispersible through their tiny, wind-dispersed seeds (Drake, 1992) and large, showy inflorescences that attract a wide variety of bird and insect species to the nectar and pollen of this keystone resource (Koch & Sahli, 2013); and the prolonged flowering periods of these varieties are highly overlapping in the field (Stacy *et al*. 2017), common garden, and greenhouse (Stacy and Powless, unpub. data). Long-distance gene flow in this species is evident in the weak to absent genetic structure observed within varieties across Hawaii Island (Stacy *et al*., 2014).

**Fig. 1.**
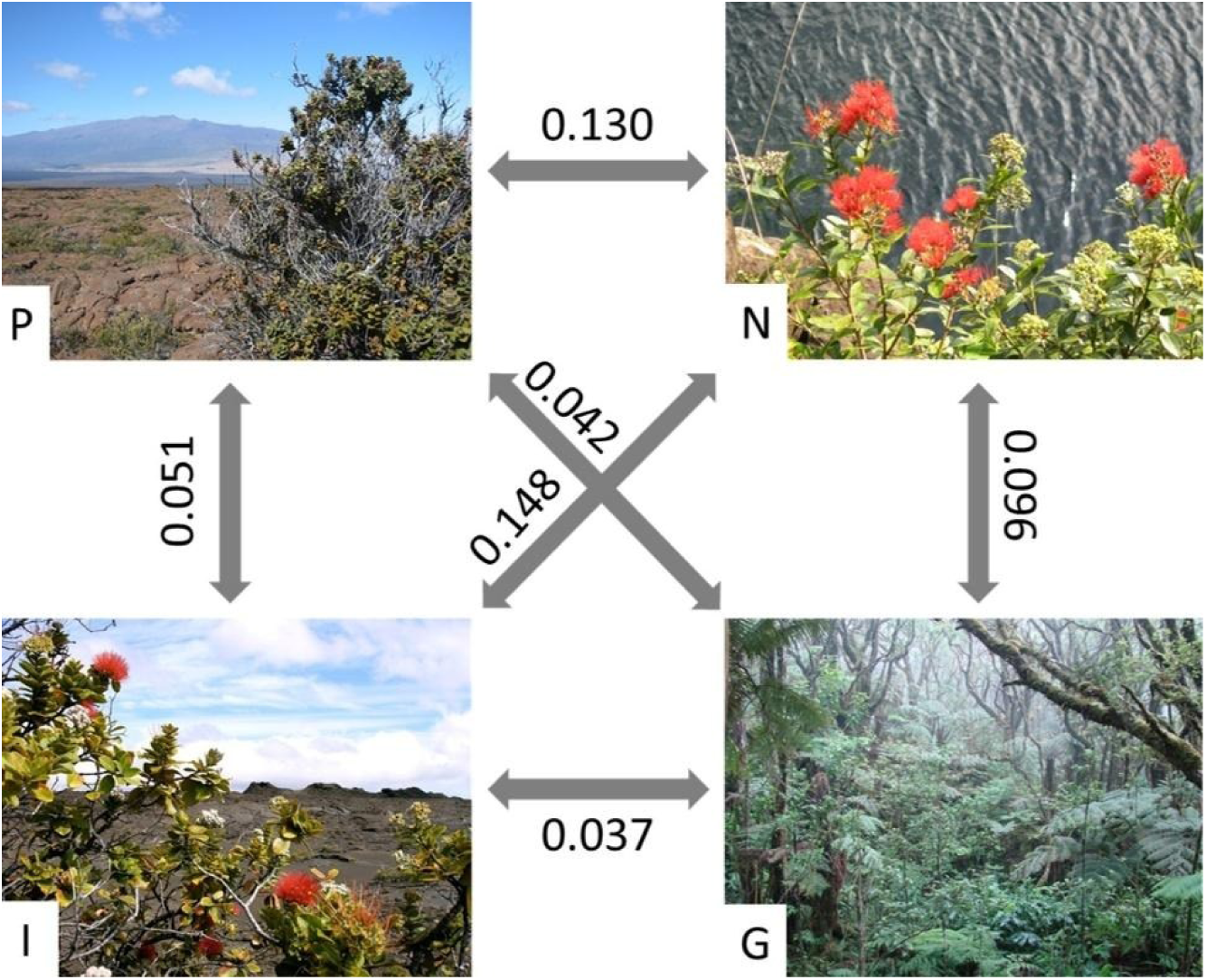
Environments of the four varieties of *M. polymorpha* on Hawaii Island and mean population-pairwise F_ST_ values based on nine nuclear microsatellite loci (Stacy *et al*. 2014). See text.

**Fig. 2.**
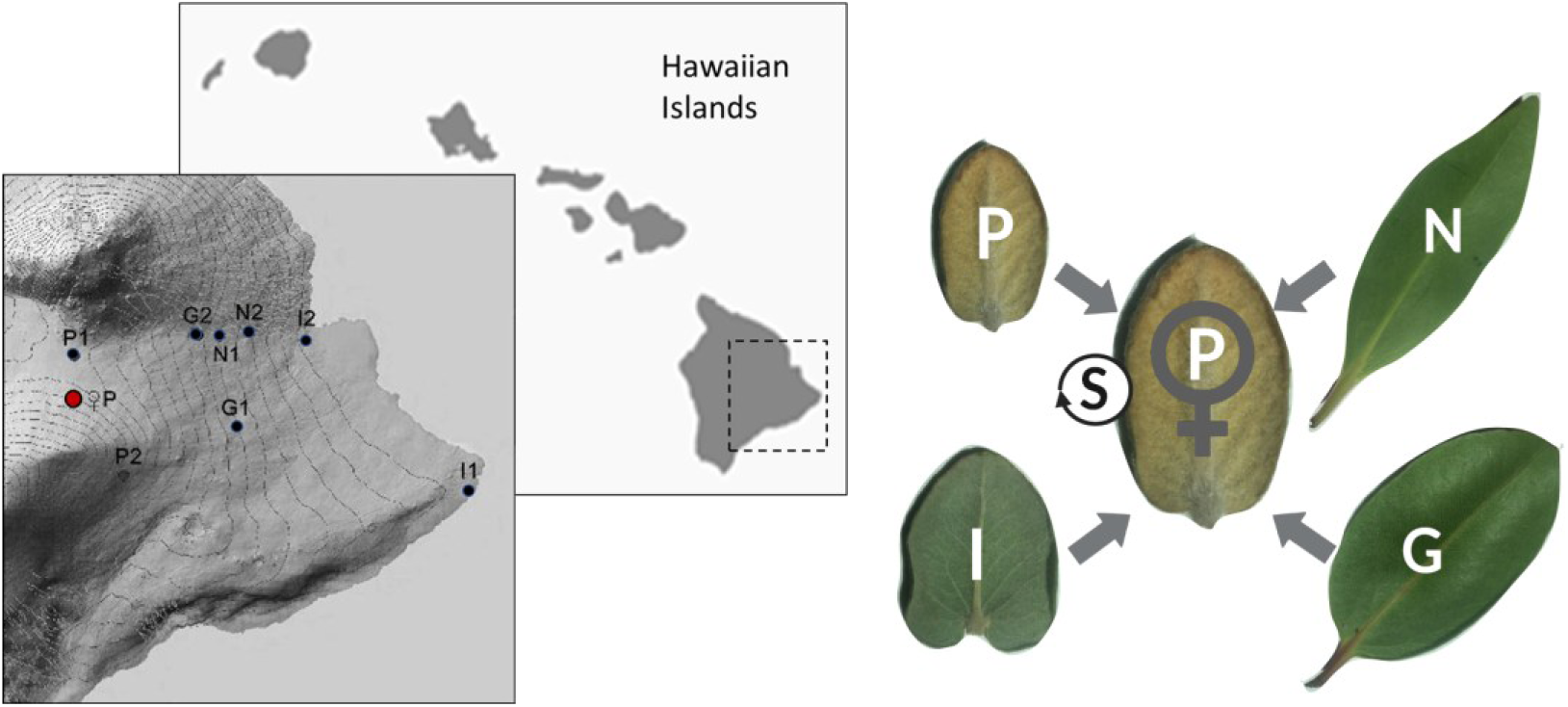
Experimental design of the field crosses. *Left:* Location of pollen-donor and maternal populations on Hawai’i Island. Pollen donors were sampled from each of two sites per variety for crossing with a single maternal population of polymorpha (♀P; red). G, I and N represent glaberrima, incana, and newellii, respectively. *Right:* Each of 21 maternal trees was pollinated with pollen from a single individual of each of the four varieties: G, I, N, and P (control). Self-pollinations (S) and monitoring of unmanipulated, open-pollinated flowers were also done on all 21 maternal trees.

Over a period of ∼14 years, we examined post-pollination barriers through fruit set of F_1_ hybrids for four of the six possible *Metrosideros* taxon pairs on Hawaii Island. The pairs include one that forms a stable hybrid zone, one that forms recurring ephemeral hybrid zones, a parapatric pair, and an allopatric pair; and flowering times in zones of sympatry are highly overlapping. Because of the weak to moderate genetic isolation among these ecologically diverged varieties and the abundance of hybrids where they co-occur, the isolating barriers observed here are expected to be among the first to evolve within this landscape-dominant tree.

## Materials and Methods

This study comprised a large number of single-pollen-donor F_1_ and within-variety (control) crosses on maternal trees of high-elevation polymorpha in the field that were monitored for rates and timing of fruit set and seed germination, after which the progeny were assessed for seedling phenotype, and survivorship, maturation rate, and fertility through ≥8 years in a greenhouse. The analyses of survivorship, maturation rate, and fertility were supplemented with “pure-taxon” (control) plants of Hawaii Island’s three other varieties and a fourth F_1_ hybrid genotype derived from companion studies (described below) for a more comprehensive analysis of early barrier evolution within *M. polymorpha*.

### Maternal and pollen-donor populations used in the field crosses

All field crosses were done on maternal trees (n=21) in the highest-elevation population of polymorpha on Mauna Loa, Hawaii Island (Fig. 2). This population was selected as a maternal population because: 1) it is likely more pure (experiencing less hybridization) than other populations as a consequence of strong selection at the edge of the species’ altitudinal range (Merondun *et al*., in prep), and 2) flowering of polymorpha occurs throughout the year, thus allowing crosses with each of the other varieties for which flowering is more seasonal. Pollen donors were selected from two populations each of incana, glaberrima, newellii and polymorpha (as a control) on east Hawaii Island (Fig. 2, Table S1). To minimize the probability of sampling hybrids, pollen donors were selected from monotypic stands (i.e., >95% of adults in a 1-km^2^ area appear to be a single variety). An exception was made for newellii, which exists in narrow populations along waterways, within glaberrima-dominated forest (Ekar *et al*., 2019). Within each population, pollen donors (n=10- 11) were chosen based on the presence of accessible (with pole-cutter), freshly dehiscing inflorescences. Collected inflorescences were pinned to damp floral foam secured within plastic containers for transport to the maternal population for same-day pollinations.

### Controlled crosses

Each of the 21 maternal trees received pollen from a single donor of each of the four varieties (Fig. 2); 11 maternal trees received pollen from the first donor population of each variety, and 10 maternal trees received pollen from the second donor population. Crosses were completed January through December 2011 and followed the methods of Stacy *et al*. (2017). Briefly, developing flower buds on the maternal trees were tagged, emasculated before anther dehiscence, and covered with a fine-mesh bag to exclude pollinators. When possible, donor inflorescences were also bagged 2-3 days before pollen collection. Flowers were pollinated 4-7 days after anthesis by brushing the stigmas directly with anthers of the donor flowers. Pollinator-exclusion bags were then replaced on the experimental inflorescences and left for 14 days. Upon bag removal, the number of developing fruits remaining was recorded as the initial fruit count (x).

### Fruit set and maturation time

For each hand-pollinated inflorescence, the number of fruit capsules retained through maturation (8-15 months post-pollination) was recorded (y) and used to calculate fruit set rate as (y/x)*100. The number of weeks to reach maturity was also recorded. *Analysis:* Fruit set rate was examined in a linear mixed effects model (LMM) with binomial error, from the lme4 library (Bates *et al*., 2015). The LMM models the probability of observing x successes in N trials, and fruit set was coded as “success, failure” beginning with a count on week 0 and ending with the final mature-fruit count. Pollen-donor variety was treated as the main fixed effect and individual maternal tree as a random effect nested within variety. The odds ratio was used to determine the probability that a flower pollinated by a donor would produce a mature fruit, and model selection was based on Akaike information criterion (AIC) values (Cavanaugh & Neath, 2019). The mean number of weeks to maturation was compared among pollen-donor types using an analogous linear mixed-effects model with a normal error structure using the same fixed and random effects. Because this was the first robust study of reproduction in polymorpha, fruit set rate and maturation time were estimated for self- and open-pollinated inflorescences also (see Supplementary Material).

### Pollen tube length and density

A separate set of outcross and self pollinations was done on five of the maternal trees in the field for the examination of pollen tube length and density following Martin (1959) and (Schmidt-Adam *et al*., 1999). Outcrosses involved pollen from only a single population per donor type (P1, N1, G1 and I1; Fig. 2, Table S1). For both outcross and self pollinations, 10 flowers per tree were pollinated 4-7 days after the onset of anthesis. On Day 7 after pollination, five styles were harvested and fixed in 70% ethanol, cleared and softened in 8 N NaOH, and stained with 0.03% decolorized aniline blue (DAB) solution for ca. 8 hr, and these steps were repeated for another five styles on Day 10. The squashed styles were examined using the UIS2 Fluorescence System in an Olympus BX51 upright microscope (Olympus Corp, Center Valley, PA). The length of the pollen-tube bundle was recorded as a proportion of style length (measured using an ocular micrometer). Pollen-tube bundle density was scored as low (< 50 tubes), medium (50-500), or high (> 500). Pollen-tube bundle length and density were scored blindly. *Analysis:* The mean percent length of style through which outcrossed pollen tubes grew 7 and 10 days after pollination was assessed in a generalized linear model with Gaussian error, in which each of incana, glaberrima, and newellii was compared to the control, polymorpha. The distribution of observed pollen tube density scores for each donor class, 7 and 10 days after pollination, was examined using multinomial logistic regression from the mlogit package (Croissant, 2020). High density was set as the reference level, as it was predicted that the control cross would yield high-density pollen tube bundles, and pollen donor = polymorpha was dummy coded as the intercept. Model selection for both analyses was based on AIC values. Pollen tube length and density of self pollinations were analyzed as above with the corresponding polymorpha outcross from the same maternal tree used as the control.

### Seed germination rate and timing

Mature fruits from the outcross on the 21 maternal trees were harvested from November 2011 through October 2012 and dried in an air-conditioned lab for 2 weeks. Four mature fruits per cross (maternal-treeXpollen-donor combination) were haphazardly sampled, and seeds were sown on sand layered on well-draining media [4 parts Sunshine #4 Pro Mix (SunGro Horticulture, Awawam, MA): 4 parts cinder: 3 parts perlite] in standard sheet propagation trays. Trays were placed in a misthouse with ∼24% sunlight and 5 s of mist every 10 min during daylight hours for 10 weeks. The number of germinants was counted weekly to determine the timing of seed germination and the maximum number of germinants per fruit. *Analysis:* The maximum number of germinants per fruit (square-root transformed) and the mean number of weeks required to reach maximum germination were analyzed in linear mixed-effects models. For each measure, the best-fit model based on AIC values included variety as the fixed effect and maternal tree as a random effect nested within variety. The seed germination rate was estimated for self-pollinated inflorescences also (see Supplementary Material).

### Seedling morphology

After 10 weeks in the misthouse, the seedling trays were transferred to a coldframe with ∼65% sunlight and overhead water 3X/day, allowed to acclimate for 2 weeks, and then thinned to six well-spaced seedlings per 5×10 cm well. Over 1,000 seedlings were then individually potted up into 58-cm^3^ pots. Seedling trays were rotated within the greenhouse monthly, and light fertilizer and pesticides were used, as needed, to maintain seedlings for another 18 months. Mortality of seedlings in both the germination trays and individual pots was negligible. At this stage (∼24 months old), 661 seedlings were haphazardly chosen from all four genotypes derived from all 21 maternal trees for the following measurements: root length, number of leaves, number of stems, main stem length, total stem length, length and width of the largest leaf, presence/absence of petioles, and abaxial (leaf) pubescence score (0 = glabrous, 1 = pubescence that is easily removed, 2 = permanent pubescence, following Stacy *et al*. (2016, 2017). *Analysis:* To compare F_1_ seedling morphology with control-seedling morphology, principal components analysis (PCA) of seven variables was done (total stem length and leaf width were removed due to pairwise correlations >0.7), followed by ANOVA of transformed scores from the three significant PC axes, with polymorpha seedlings used as reference group.

### Additional companion-study plants used in analyses of F_1_ fitness

As the field crosses produced seedlings of only one parental taxon (polymorpha), additional “control” plants of all four Hawaii Island *Metrosideros* taxa were leveraged from other studies for comparison of survivorship, maturation rate and fertility (see below). Plants from a fourth Hawaii Island F_1_ genotype, glaberrimaXincana (GI), were also used. The additional glaberrima, incana and GI F_1_ hybrid plants derived from controlled crosses in an ephemeral GI hybrid zone on east Hawaii Island (Stacy *et al*., 2016, 2017), and all newellii plants as well as additional incana and polymorpha plants were derived from open-pollinated fruits collected from maternal trees also on east Hawaii Island (Ekar *et al*., 2019; Merondun *et al*., in prep). All plants derived from companion studies were grown from seeds at the same facility and following the same protocols used in the polymorpha-centered study, and all were monitored for survivorship, maturation rate, and fertility through age ≥8 years following the same protocols used for the polymorpha-derived plants. Plants from Ekar *et al*. (2019) and Merondun *et al*. (in prep) were grown from seed contemporaneously with plants of the focal F_1_ genotypes and polymorpha controls, while plants from Stacy *et al*. (2016, 2017) were started from seed (and thus measures were recorded) roughly 5 years earlier.

### Survivorship and maturation rate of four varieties and four F_1_ genotypes in the greenhouse

Following measurements at 2 years of age, a subset of seedlings was individually potted in 1,400-cm^3^ pots and maintained in the greenhouse with light fertilizer, pesticides, and potting up, as needed, through age 8 (years). Pots were rotated every 2 months until they became too heavy to move, and the date of death was recorded for any plant that died. Beyond that stage, all surviving individuals of all genotypes except newelliiXpolymorpha (NP), which showed high survivorship and flowering by age 8 years, were retained for continued monitoring of maturation rate and fertility through age 12 (years). *Analysis:* The percentage of potted-up plants that survived to age 8 and that survived and flowered by age 8 were plotted. Age (years) at first flowering (i.e., maturation rate) through 12 years was compared among the four varieties using ANOVA, followed by Tukey’s pairwise comparisons after homogeneity of variances was confirmed with Levene’s Test. Maturation rates of the four F_1_ genotypes were then compared to the maturation rates of their respective parental varieties through four Kruskal-Wallis tests and Dunn’s posthoc comparisons due to non-homogeneous variances.

### Fertility of four varieties and four F_1_ genotypes in the greenhouse

Upon flowering, trees of the four parental taxa were used in within- and between-variety crosses, and the F_1_ trees were used in reciprocal, controlled F_2_ and/or backcross pollinations with trees of the parental taxa. Pollen stainability with lacto-phenol cotton blue (Kearns & Inouye, 1993) was examined when additional pollen was available. The controlled pollinations followed the protocol for outcrosses in the field, except that not all crosses were independent due to limited flowering of some genotypes, and pollinator-exclusion bags were not used in the pollinator-free greenhouse. The rate of fruit set and duration of the fruit-maturation period were recorded for each outcrossed inflorescence following the methods used in the field. A second set of crosses was done for the examination of pollen tube length and density at Day 7 post-pollination, following the field methods, except that density was assessed separately at the distal, middle, and proximal thirds of each style. Pollen tubes were not examined at Day 10, as all tube bundles had reached the base of the style by Day 7 in the low-elevation greenhouse. *Analysis: Fruit set rate and maturation time:* For each measure, because of the unbalanced sampling across cross types and genotypes, the data were pooled into two groups based on maternal-tree class (parental variety or F_1_) followed by a Wilcoxon rank sum test of differences between groups. *Pollen tube growth and pollen stainability:* Because pollen-tube bundles had reached the base of the style in all 111 pollen-tube crosses, bundle lengths are not reported. Rather, variation in style lengths among taxa was examined using ANOVA, followed by Tukey’s pairwise comparisons. Again, because of uneven sampling, both pollen-tube-bundle density and pollen stainability were analyzed through Wilcoxon rank sum tests with crosses pooled by maternal-tree class. For the pollen-tube data, this same test was then repeated with crosses pooled analogously by pollen-donor class.

All analyses were executed in R 4.3.0 (“R core team,” 2023).

## RESULTS

### Fruit set, seed germination, and pollen-tube growth of outcrosses on polymorpha

*Rate and duration of fruit set—*A mean of 28 flowers were pollinated per maternal-treeXpollen-donor combination (2,634 total) on 21 maternal trees of polymorpha, and 74.6% of the outcrossed flowers produced mature fruits. The proportion of hand-pollinated flowers that set mature fruits was lower for newellii pollen donors compared to the control (z=-2.76, p=0.006; Fig. 3; Table S2); flowers from these crosses were 48% as likely as the control flowers to mature fruits. Fruit set of crosses with glaberrima or incana did not differ from control fruit set. Fruit maturation time averaged 45.1 weeks (ca. 11 months) and varied considerably among maternal trees (Fig. S1), but did not differ between any F_1_ cross (glaberrima: 43.83 +/- 1.16 (SE) weeks, incana: 44.86 +/-1.40 weeks, newellii: 46.03 +/- 0.70 weeks) and the control (45.75 weeks +/- 1.64), which was intermediate (Table S3).

**Fig. 3.**
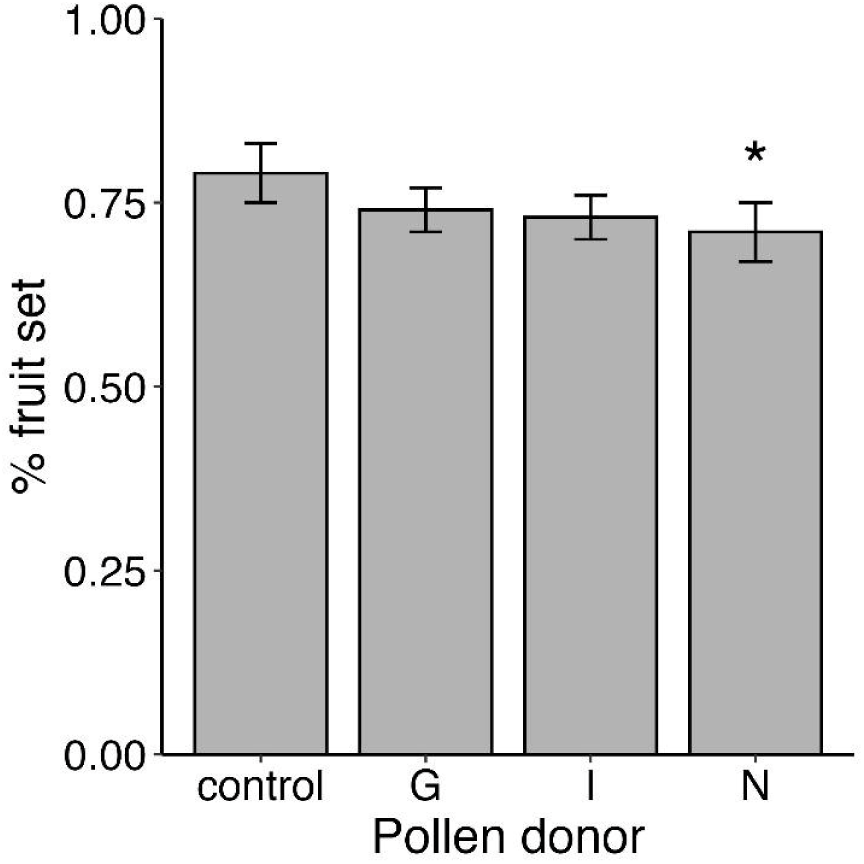
Percentage (mean ± 1 SE) of hand-pollinated flowers on 21 maternal trees of polymorpha that resulted in mature fruits for each of four pollen donor varieties; control = polymorpha. Taxon codes are as in Fig. 2; * indicates significant difference from the control.

*Rate and timing of seed germination—*The maximum number of germinants per outcrossed fruit ranged from 1 to 415. Germination rates ranged from just 52.2 +/- 7.41 (SE) per fruit for incana to 95.7 +/- 10.7 for glaberrima, yet none of the F_1_ crosses differed from the control, which was intermediate (79.4 +/- 10.7 germinants/fruit); Fig. S2, Tables S4 and S5).

Time to maximum germination ranged from 3.15 +/- 0.19 (SE) weeks to 4.90 +/- 0.24 weeks for incana and newellii, respectively, and again none of the F_1_ crosses differed from the control, which was again intermediate (3.79 +/- 0.21 weeks; Fig. S3).

*Pollen tube growth—*There was a nearly significant reduction in pollen tube length in flowers pollinated by incana compared to the control cross (Day 7: t=2.1, P=0.052; Day 10: t=- 2.06, P=0.056; Fig. 4). Pollen tubes sired by glaberrima grew to 32.94 +/- 2.90% (SE) of style length on Day 7 and 49.03 +/- 7.39% on Day 10 and did not differ from the control (Day 7: 32.60 +/- 2.31%; Day 10: 49.20 +/- 3.22%); newellii-derived pollen tubes also did not differ in length from the control cross at either time point, but trended longer than the controls by Day 10 (56.07 +/- 7.04% of style length; Fig. 4). Pollen-tube bundle density was robust (medium or high) in most crosses. The exception was that of glaberrima-pollinated flowers, for which density was significantly lower than that of the controls at both 7 and 10 days after pollination, with the difference narrowing over time (Day 7: 77.0% of control, t=2.46, P=0.014; Day 10: 81% of control, t=2.01, P=0.045; Fig. 5).

**Fig. 4.**
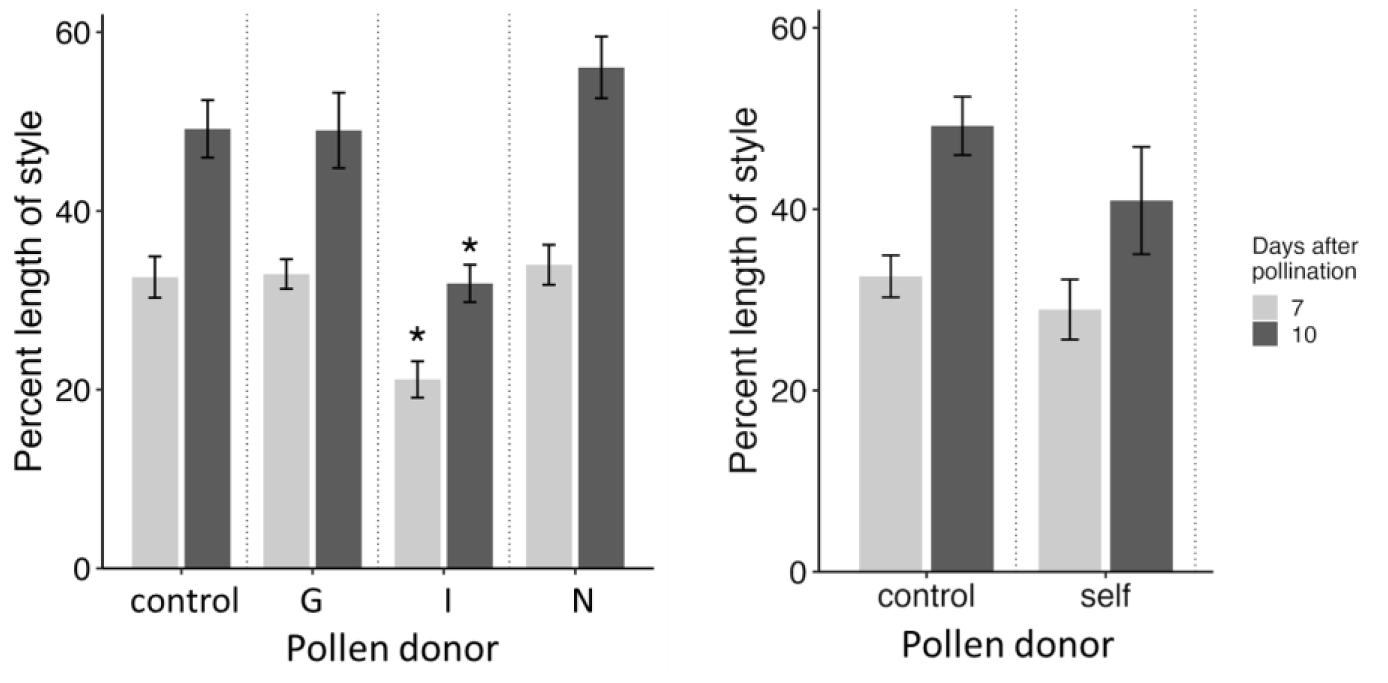
Mean (+/- 1 SE) percent length of style through which pollen tubes grew 7 and 10 days after pollinations between each of four pollen-donor varieties and five maternal trees of polymorpha. *Left:* Outcrosses used pollen from four varieties; control=polymorpha. Taxon codes are as in Fig. 2. *Right:* Self pollinations were compared with outcrossed (control) pollinations on the same maternal trees. * indicates near-significant difference from the control.

**Fig. 5.**
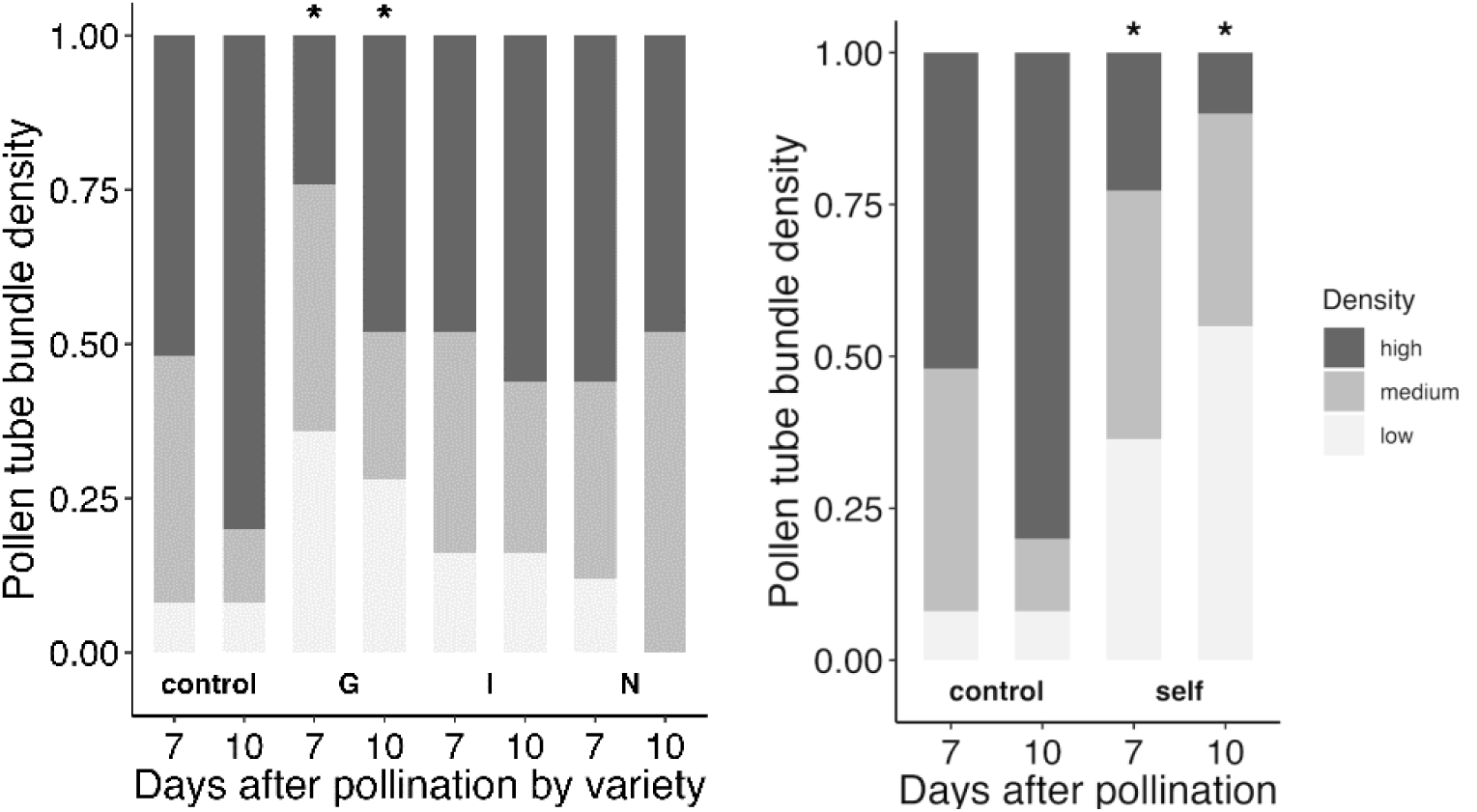
Distribution of pollen-tube bundle density scores (low, medium, high) observed 7 and 10 days after pollination of five maternal trees of polymorpha. *Left:* Outcrosses used pollen from four varieties; control=polymorpha. Taxon codes are as in Fig. 2. *Right:* Self pollinations were compared with outcrossed (control) pollinations on the same maternal trees. * indicates significant difference from the control.

#### Fruit set, seed germination, and pollen-tube growth of self and open pollinations on polymorpha

Fruit set was reduced for self-pollinated flowers, but not for open-pollinated flowers, while seed germination was reduced for both cross types, relative to controls (see Supplementary Material). On Days 7 and 10, pollen tubes in self-pollinated styles grew on average 32.60% +/- 6.69 SE and 40.96% +/- 13.28, respectively, of the style length, which was not different from the control values (above; Fig. 4). Pollen-tube bundle density, however, was lower in self-pollinated styles than control styles at Day 7 (76.3% of control; t=2.30, P=0.021) with a larger difference at Day 10 (57.0% of control; t=3.75, P<0.001; Fig. 5).

#### Morphology of outcrossed seedlings

The PCA of seven morphological traits of all 2-year-old outcrossed seedlings pooled yielded three significant axes. Mean scores for PC1 (35.0%), which represented plant size, were greater for incana-polymorpha (IP) (t=2.33, P=0.02) and especially newellii-polymorpha (NP) (t=6.25, P<0.001) F_1_ hybrids compared to the control (polymorpha) seedlings; while glaberrima-polymorpha (GP) F_1_ seedlings trended smaller than the controls (t=- 1.84, P=0.065; Fig. 6). Mean scores for PC2 (19.1%; leaf length vs. leaf number) and PC3 (14.4%; pubescence) also differed between F_1_ and control seedlings (Fig. S4). Excluding pubescence score, the means of the individual measures were generally highest for the NP F_1_s, while all above-ground measures except leaf size (i.e., leaf length and width and petiole length) were lowest for the GP F_1_s (Fig. 7).

**Fig. 6.**
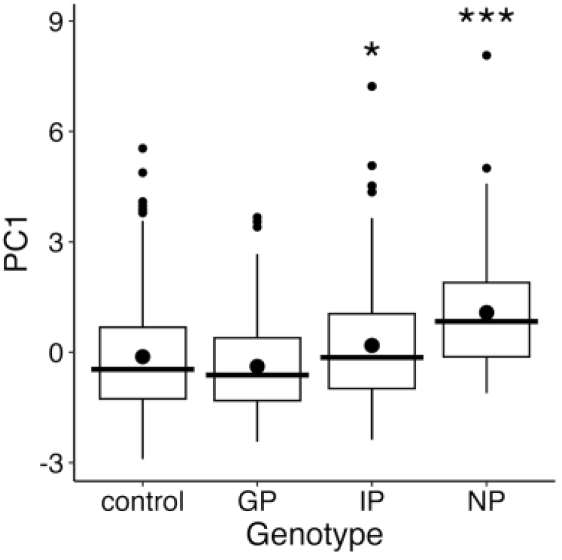
Boxplots of variation in morphology (PC1 scores) among 2-year-old seedings representing four genotypes produced through controlled crosses on 21 maternal trees of polymorpha. Genotypes: control=polymorpha, GP=glaberrimaXpolymorpha, IP=incanaXpolymorpha, NP=newelliiXpolymorpha. Circles indicate means; lines indicate medians. * indicates significant difference from the control.

**Fig. 7.**
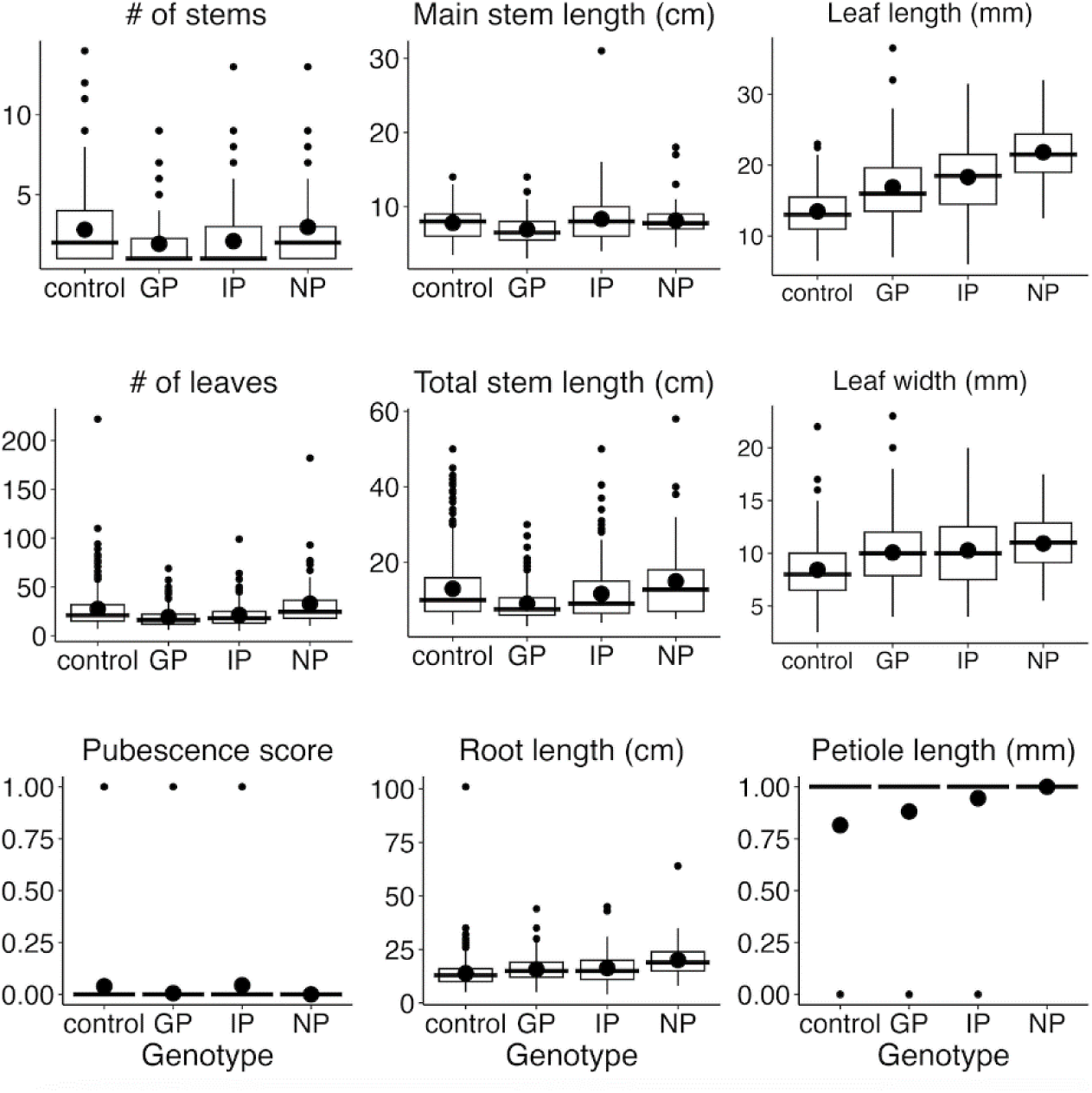
Boxplots of variation in nine morphological traits among 2-year-old seedings representing four genotypes produced through controlled crosses on 21 maternal trees of polymorpha. Genotypes are as in Fig. 6. Closed circles indicate means; lines indicate medians. * indicates significant difference from the control.

#### Survivorship, flowering, and maturation rates of four varieties and four F_1_ genotypes

Survivorship between ages 2 and 8 years of potted-up plants of the parental varieties was high for the two successional varieties, incana and glaberrima (92.7 and 95.5%, respectively), and slightly lower for the riparian variety (newellii; 76.2%) and high-elevation variety (polymorpha; 72.0%; Fig. 8). F_1_ survivorship between ages 2 and 8 varied more than two-fold among genotypes, from 41.3% for GP to >90% for GI, IP, and NP (Fig. 8). Notably, survivorship of the GP and NP F_1_ hybrids fell well below (50%) and above (14%) the averaged survivorship of their respective parental varieties (Fig. 8). Beyond age 8, survivorship of the retained GP F_1_s continued to decline. As of June 2025 (age 13 years), only seven (15.2%) of the original 46 GP F_1_s had survived, whereas all of the retained GI, IP and NP F_1_s survived the same 5-year period or to the mature fruit-set stage, whichever came first.

**Fig. 8.**
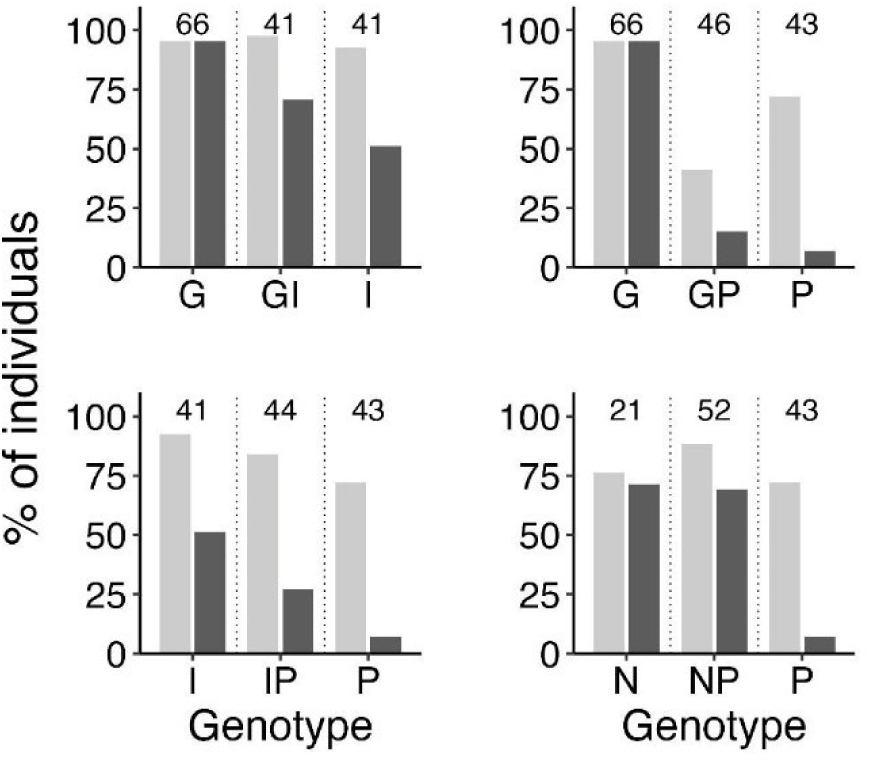
Survivorship and flowering of four pairs of varieties of *M. polymorpha* from Hawaii Island and their F_1_ hybrids in a greenhouse. Shown are the percentage of plants potted up at age 2 (years) post seed-germination that survived to age 8 (years; light bars), and the percentage of potted-up plants that survived *and* flowered by age 8 (years; dark bars). Genotypes are as in Fig. 6. Sample sizes for the eight genotypes are shown along the tops of the panels.

The percentage of ‘pure-variety’ plants potted up at 2 years that flowered by 8 years ranged from 7% for polymorpha to 95.5% for glaberrima, with incana (51.2%) and newellii (71.4%) in between. The percentage of potted-up plants that survived and flowered by 8 years varied among the F_1_ hybrid genotypes as well, from 15.2% and 27.3% for GP and IP, respectively, to 70% for GI and NP; all four F_1_ genotypes fell between their parental varieties in this measure (Fig. 8).

The age at first flowering (maturation rate) through 12 years differed among the four varieties (F_3_=36.1, P<0.001). Glaberrima matured more quickly than all others (median 4 years) but had the broadest range of maturation times (1 to 8 years); polymorpha was the slowest to mature (8 years), while incana and newellii (5 and 6 years, respectively) were intermediate (Fig. 9, Fig. S5). Maturation rates of three of the four F_1_ genotypes were slightly slower than intermediate between their parental taxa and not different from that of the slowest parent; NP F_1_ hybrids, in contrast, flowered significantly more quickly (median: 5 years) than either parent (6 and 8 years; H=18.0, df=2, P<0.0001; Fig. 9, Fig. S5).

**Fig. 9.**
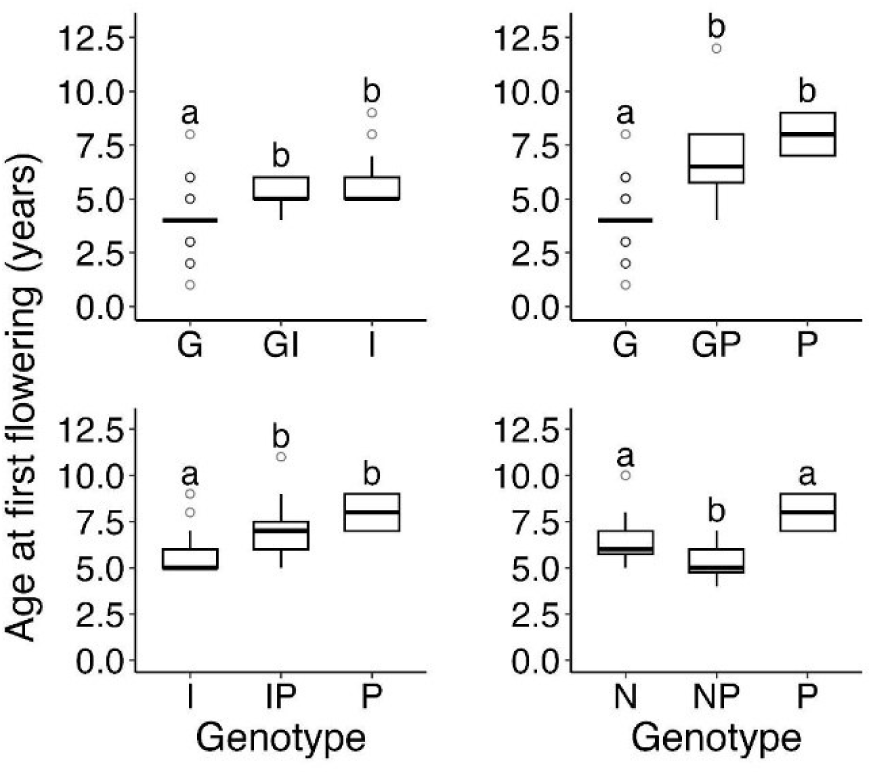
Boxplots for age at first flowering of four pairs of varieties of *M. polymorpha* from Hawaii Island and their F_1_ hybrids in a greenhouse. Genotypes are as in Fig. 6; sample sizes are as in Fig. 7. Shared superscripts indicate no significant difference.

#### Fertility of four varieties and four F_1_ genotypes

*Fruit set and fruit maturation time*—Fruit set rate of 128 pooled outcrosses on pooled F_1_ maternal trees was 44% higher (median: 81.8%) on average than that of the 132 outcrosses on pooled maternal trees of the four parental varieties (56.8%; W=10,114, P<0.01). Given the low fruit set rate of maternal trees of polymorpha in the greenhouse, fruit set rates of the pooled maternal F_1_s and pooled parental varieties were compared again with polymorpha removed, but fruit set of F_1_ trees was still higher (W=8,826.5, P=0.01). Median fruit set of all three polymorpha F_1_ hybrids (i.e., GP, IP, NP) exceeded that of both parents (Fig. S6), whereas fruit set rate of GI F_1_ trees was intermediate to that of the parental taxa. Fruit maturation time of maternal trees (with pollen donors pooled) was highly variable within genotypes, and ranged from just 22.1 weeks (median) for incana to 31.6 weeks for newellii, 32.9 weeks for glaberrima, and 34.4 weeks for polymorpha. The fruit maturation time of pooled maternal F_1_ hybrid trees did not differ from that of the pooled varieties, as F_1_s were generally intermediate to their parental taxa (Fig. S7).

*Pollen stainability*—A mean of 369 stained pollen grains (range: 109-1,131) were examined per tree from 103 greenhouse trees (mean=12.9 trees/genotype; range: 1-33). The mean percentage of pollen grains that appeared normal was high and not different between the pooled F_1_ genotypes (median: 90.2%; range: 54.9-98.7%) and the pooled varieties (93.6; 51.3- 98.9%; Fig. S8).

##### Style lengths and pollen-tube density

Mean style length was greater for newellii (22.3 +/- 1.4 mm (SE), n=10) than for incana (18.2 +/- 0.7 mm, n=17; F_2,30_=4.2, P=0.025), with styles of glaberrima intermediate (19.4 +/- 1.2 mm, n=6). Styles from the single individual of polymorpha used as a maternal tree in the pollen-tube crosses were long (24.1 mm). Mean style lengths of the F_1_ genotypes were intermediate to their parental means, except GP, which trended lower (no statistics due to uneven sampling). There was no difference in pollen-tube bundle density (at any position within the style) between pooled parental varieties and pooled F_1_ trees when grouped by either maternal-tree type or pollen-donor type. However, the observed variation in bundle densities may suggest significant variation with additional sampling (e.g., low and high bundle densities in maternal trees of GP and NP F_1_ hybrids, respectively; Fig. S9).

## DISCUSSION

We characterized post-pollination, pre-zygotic and early post-zygotic barriers among the four varieties of *M*. *polymorpha* that specialize on different environments within the large, continuous *Metrosideros* forest on Hawai’i Island and found four contrasting patterns and strengths of early barrier evolution. Because of the concurrent flowering and extensive pollen flow in this species, ecogeographic isolation and immigrant inviability are not expected to isolate these varieties in the vast regions where they co-occur. Instead, isolation of co-occurring varieties appears to manifest through a mix of predominantly postzygotic mechanisms.

### Four patterns in four F_1_ crosses in M. polymorpha

This study, coupled with a previous study of isolating barriers between the successional varieties, incana and glaberrima (Stacy *et al*., 2017), revealed four contrasting patterns and strengths of post-pollination prezygotic and early postzygotic isolation among four taxon pairs, along with contrasting patterns of F_1_ hybrid vigor. These results support the finding that the evolution of isolating barriers in plants varies among taxon pairs (Moyle *et al*., 2004; Lowry *et al*., 2008; Baack *et al*., 2015), but here, demonstrates that variability at the early stages of diversification, and in trees. The nature of the barriers observed between varieties of *M. polymorpha* on Hawaii Island appears to be shaped by genotype and the persistence of sympatry on this volcanically active island.

### Reduced fruit set followed by hybrid vigor in allopatric newellii-polymorpha

The one-way crosses between newellii and polymorpha were marked by significantly reduced fruit set, followed by hybrid vigor in F_1_ plants. The NP F_1_ seedlings were the most robust of the five genotypes measured, and qualitative comparisons of the NP seedling measurements with those of comparably-aged newellii seedlings from another study (Ekar *et al*., 2019) suggest hybrid vigor of NP F_1_s at the seedling stage. These F_1_ plants subsequently reached the flowering stage more quickly than either parental taxon, and all other F_1_ measures trended higher than those of both parents.

Newellii and polymorpha are the most strongly isolated *Metrosideros* taxon pair on Hawaii Island (F_ST_ = 0.13) (Stacy *et al*., 2014) and the only pair that can be considered allopatric. Within the continuous *Metrosideros*-dominated forest on east Hawaii Island, which extends from sea level in some areas to over 2400 m, newellii and polymorpha have the two most restricted distributions, currently and historically (Izuno *et al*., 2017). Newellii is particularly constrained, occurring in narrow populations <915 m along waterways, where populations separated by just a few km are significantly differentiated by drift (Stacy *et al*., 2014). Given their distributions, significant divergence between newellii and polymorpha through both selection and drift is expected, and gene exchange between them - either current or historical - is unlikely. The vigorous pollen-tube growth in the newellii-polymorpha crosses and trend toward longer pollen tubes relative to controls are consistent with an absence of pollen discrimination between closely related allopatric taxa, as expected if pollen discrimination arises predominantly through reinforcing selection in sympatry (Dobzhansky, 1937; Coyne & Orr, 2004), and may also be associated with long pollen tubes in newellii, which, along with polymorpha (Stacy & Johnson, 2021), has long styles (this study). The lower fruit set of polymorpha flowers pollinated with newellii pollen, however, indicates the presence of a partial, but significant, late-prezygotic or early postzygotic barrier, possibly associated with problems arising during hybrid embryo development (Nakamura, 1986). Lastly, the hybrid vigor of NP F_1_s observed at multiple stages is consistent with a release from inbreeding depression within the parental taxa due to the accumulation of contrasting genetic loads in isolated populations (Lohr & Haag, 2015).

### Mismatched pollen tubes and styles in incana-polymorpha

The one-way crosses between incana and polymorpha (F_ST_ = 0.051) (Stacy *et al*., 2014) were marked by shorter pollen tubes (P = 0.05) relative to the control crosses followed by a non-significant reduction in the number of germinants per fruit. No other barriers were observed, and the measures of growth and fertility of the IP F_1_ hybrids were either intermediate to, or greater than, those of the parental taxa. These results suggest that a partial one-way prezygotic barrier has formed between these varieties and that intrinsic postzygotic barriers are absent through the F_1_ fruit-set stage.

The weak one-way prezygotic barrier observed between incana and polymorpha is likely a byproduct of their contrasting distributions along the 2400-m elevation gradient of *Metrosideros* forest on east Hawaii Island. Incana and polymorpha share the trait of leaf pubescence and are the first trees to colonize fresh lava flows - incana at lower elevations and polymorpha at higher elevations. Across this long elevation gradient, pollinator communities vary, with birds serving as primary pollinators at high elevations (Hart *et al*., 2011), and this shift was suggested as a likely driver of divergence in floral style length between incana and polymorpha in natural populations (Carpenter, 1976) and in a common garden (Stacy & Johnson, 2021). Differences in style length can affect fertilization rates and effect prezygotic isolation (Gore *et al*., 1990; Sorensson & Brewbaker, 1994; Tiffin *et al*., 2001; Larcombe *et al*., 2016a).

The heritable difference in style length observed between incana and polymorpha, coupled with the observation of shorter pollen tubes in one-way crosses between them, suggest that differential adaptation to contrasting pollinator communities at low and high elevations on the high Island of Hawaii may be leading indirectly to partial, one-way isolation between these varieties. The corresponding non-significant reduction in seed germination in 21 independent crosses, however, suggests that this barrier is very weak.

### F_1_ hybrid breakdown and nascent reinforcement in glaberrima-polymorpha

The one-way crosses between glaberrima and polymorpha were marked by reduced pollen-tube density, high seed germination, and striking F_1_ hybrid weakness at multiple stages. While seedlings of polymorpha and glaberrima have the slowest and fastest above-ground growth rates, respectively, observed within Hawaii-Island *Metrosideros* under ambient greenhouse conditions (current study, Stacy *et al*., 2017), the 2-year-old GP F_1_ seedlings were equal to polymorpha in all measures, except leaf size. At later stages, the GP F_1_ hybrids showed severely compromised fitness through both reduced survivorship and fertility. Specifically, only 19 of 46 potted-up 2- year-old GP F_1_s survived to age 8 years. While the percentage of surviving trees that flowered (44%) was roughly intermediate to that of polymorpha (7%) and glaberrima (95.5%), flowering of all but one of these GP F_1_s was minimal (i.e., <5 inflorescences total over all flowering periods). Of the 18 outcrosses that were possible on GP maternal trees: 14 (77.8%) were restricted to just two sibs. Only three of the eight pollinated GPs, primarily the two sibs, were capable of setting fruit; four of the five others that received outcross pollen either died (n=1) or sacrificed the pollinated branch (n=3) shortly after pollination. Only 10 crosses were possible with GP F_1_s as pollen donors, again predominantly (80%) with the two sibs. By age 13 years, survivorship of GP F_1_s had dropped to 15.2%. In sum, just one (< 5%) of 21 independent glaberrima-polymorpha crosses in the field yielded F_1_ hybrids with some degree of fitness as measured in the greenhouse. This prolonged period of GP F_1_ fitness decline over 13 years following high seed germination rates supports the contention that hybrid incompatibilities in trees can manifest over a prolonged period (Lopez *et al*., 2000; Barbour *et al*., 2006; Levin, 2012; Costa e Silva *et al*., 2012; Lindtke *et al*., 2014; Larcombe *et al*., 2016a,b).

A companion study of these same trees revealed that the GP F_1_ hybrids were also unique among the four F_1_ genotypes in having lower-than-expected levels of tannins, phenols, and lignins, while the three other F_1_ genotypes were intermediate to their parental taxa in these measures (inferred through imaging spectroscopy of leaves; Seeley *et al*., 2023). The poor performance of GP F_1_ hybrids in the current study indicates significant intrinsic incompatibilities between glaberrima and polymorpha (Fishman & Sweigart, 2018), and the reduction in tannins, phenols and lignins in GP F_1_s may suggest incompatibilities associated with the shikimate pathway or downstream aromatic amino acid pathways (Vogt, 2010; Maeda & Dudareva, 2012). Disruption of the shikimate or downstream pathways may be sufficient to explain both the slower growth of GP F_1_ seedlings and their high mortality through age 13 years (Gordon *et al*., 2022).

The significant reduction in pollen-tube density in the glaberrima-polymorpha crosses in the field, followed by high seed germination, suggests the nascent effects of reinforcing selection against the formation of unfit hybrids in the only taxon pair that forms a stable hybrid zone (Dobzhansky, 1937; Coyne & Orr, 1989; Hoskin *et al*., 2005). Pollen-tube growth was slowed in these crosses, but not to a degree that prevented fertilization (i.e., pollen tubes reach the ovary before abscission of the style). Slowed growth of glaberrima-derived pollen tubes may be associated with the discrimination of self pollen observed uniquely in polymorpha styles (i.e., no self-pollen discrimination was observed in glaberrima or incana styles)(Stacy *et al*., 2017), as mechanisms that control the compatibility of both self and interspecific pollen occur in other plants (Wang & Filatov, 2023). Notably, glaberrima and polymorpha are the only taxon pair that forms a large, persistent hybrid zone on Hawaii Island. This hybrid zone, which comprises mixed phenotypes, spans upper-middle elevations and is stable for long periods due to the late-successional status of glaberrima and the lack of successional overturn of polymorpha (i.e., this variety is mono-dominant on lava flows of all ages at high elevations). These varieties have the second lowest genetic distance (F_ST_ = 0.042) among Hawaii-Island taxon pairs (Stacy *et al*., 2014), consistent with introgression. This large, persistent hybrid zone provides the sustained opportunities for hybridization that, along with the low F_1_ fitness, should promote the evolution of reinforcement (Coyne & Orr, 2004; Ortiz-Barrientos *et al*., 2009; Hopkins, 2013).

### Segregating incompatibilities in successional varieties, glaberrima and incana

The current study examined the successional taxon pair, incana and glaberrima, at three F_1_ stages only, while additional measures of cross-fertility in this pair are reported elsewhere (Stacy *et al*., 2017).

Combined, these studies revealed no post-pollination prezygotic barrier, followed by significant hybrid vigor at the F_1_ seed-germination stage (Stacy *et al*., 2017), and F_1_ plants that were largely intermediate to the parental taxa in years-2-8 survivorship, maturation rate, and fruit set (current study). These latter measures followed significant stunting (11.3%) of F_1_ plants at the early seedling stage, however, that typically led to death within 2 years. The stunted phenotype was strikingly similar to the autoimmune hybrid phenotype described in *Arabidopsis thaliana* (i.e., hybrid necrosis) (Bomblies & Weigel, 2007; Li & Weigel, 2021) and to the abnormal, stunted F_1_ seedlings of *Eucalyptus grandis* that over-expressed immune-system genes (Fuchs *et al*., 2015), but this was not recognized at the time of publication (see Stacy *et al*., 2017). The stunted phenotype was observed at a higher frequency (20%) in 575 seedlings produced through crosses with hybrid maternal trees in a natural glaberrima-incana hybrid zone (Stacy *et al*., 2017).

The modest, predominantly postzygotic, isolation observed between the successional varieties despite significant ecological divergence may be due to their extended history of alternating periods of disruptive selection and hybridization. The significant stunting observed in GI-F_1_ and advanced-generation hybrid seedling populations indicates intrinsic incompatibilities between incana and glaberrima (Fishman & Sweigart, 2018); yet the modest frequencies of the stunted phenotype indicate that the incompatible alleles are far from fixed in the parental populations. Interestingly, severe stunting was observed also within the parental taxa (18.9% of 739 glaberrima seedlings and 8.5% of 341 incana seedlings)(Stacy *et al*., 2017), indicating the segregation of incompatible alleles within these varieties as well. Unlike most Hawaiian *Metrosideros* taxa, which are endemic to individual islands or even volcanoes, incana and glaberrima occur in large populations nearly and fully archipelago-wide, respectively. On volcanically active east Hawaii Island, incana and glaberrima dominate the mosaic of new and old lava flows, respectively, below roughly 1200 m (Mueller-Dombois, 1983; Mueller-Dombois, 1987; Drake & Mueller-Dombois, 1993; Kitayama *et al*., 1995), and form ephemeral hybrid zones on intermediate-aged flows that may persist for several generations (Kitayama *et al*., 1997; Stacy *et al*., 2016, 2017). This cycle of disruptive selection followed by succession and hybrid-zone formation repeats with every volcanic eruption. Consistent with their recurring hybridization that likely dates back to Oahu at least (>3 MYA), incana and glaberrima are the most weakly isolated *Metrosideros* taxon pair on east Hawaii Island (F_ST_ = 0.037) (Stacy *et al*., 2014). The observation of stunted seedlings in all genotypes suggests the segregation of incompatibility alleles within both varieties, leading to significant incompatibilities in their hybrids (Cutter, 2012). It may be that differential local adaptation of incana and glaberrima on new and old lava flows promotes the accumulation of hybrid incompatibilities even with gene flow (Bank *et al*., 2012), but recurring selection on incompatible allele combinations in ephemeral hybrid zones, where disruptive selection is weak, prevents the fixation of these alleles within varieties. The purging of incompatible alleles in hybrid zones is consistent with the decreasing genetic distance - and thus reduced isolation - between incana and glaberrima observed with increasing island age (and, hence, time on extinct volcanoes; Stacy & Sakishima, 2019).

### Importance of post-zygotic barriers early in tree speciation

This work, coupled with studies of other species-rich tree groups, suggests that postzygotic isolating barriers are more important than prezygotic barriers at the early stages of speciation with gene flow in trees. With one exception, the barriers identified through the F_1_-fertility (fruit set) stage in this study were weak or moderate, and hybrid vigor was apparent in all crosses. Such observations are consistent with expectations at the early stages of speciation and the supposition that we are looking at the first barriers to arise in this system. In contrast to recent reviews that highlight the importance of floral character divergence at the early stages of plant speciation (Lowry *et al*., 2008), only one of four crosses revealed a weak, incipient barrier associated with floral character divergence.

These incana-polymorpha crosses were done between populations at or close to the opposite ends of the species’ elevation range, and it is possible that style lengths of these varieties are not different at the upper and lower edges of the distributions of incana and polymorpha, respectively, where hybridization is more likely. The three other crosses revealed intrinsic barriers at the stages of F_1_ fruit set and F_1_ seedling growth, with the strongest barrier apparent in reduced F_1_ sapling survivorship to adulthood and subsequent flowering. Notably, F_1_ seed germination was not reduced in any cross. These results mirror those from other trees in which predominant outcrossing, long-distance gene flow, and insufficient prezygotic barriers lead to a primary role for postzygotic mechanisms - largely beyond the seed-germination stage (Barbour *et al*., 2006; Lepais *et al*., 2013) - in the isolation of sympatric species (Lopez *et al*., 2000; Costa e Silva *et al*., 2012; Larcombe *et al*., 2014, 2016a,b; Lindtke *et al*., 2014).

The observation of significant hybrid incompatibilities in two sympatric/parapatric taxon pairs but only weak accompanying prezygotic barriers is consistent with the classic speciation model through which postzygotic barriers arise first followed by the emergence of prezygotic barriers through reinforcement (Dobzhansky, 1937; Coyne & Orr, 2004; Gavrilets, 2004). For populations that hybridize, such as closely related trees, simulations under a broad range of conditions indicate that even weak postzygotic barriers are more effective than strong prezygotic barriers in isolating populations and are thus more important at the early stages of speciation (Irwin, 2020). The finding of a nascent, post-pollination prezygotic barrier between the taxa that form a stable hybrid zone but not between the successional varieties that track active volcanoes is consistent with the prediction that the evolution of reinforcement requires both reduced F_1_ fitness and persistent opportunities for hybridization (Hopkins, 2013) in stable environments.

## Conclusions

This study examined post-pollination isolating barriers through F_1_ fertility among four ecologically diverged varieties of the highly dispersible, landscape-dominant species, *M. polymorpha*, on Hawaii Island. The results reveal a complex picture of barrier evolution likely influenced by genotype and the persistence of sympatry on this volcanically active island and add to growing evidence for the importance of postzygotic barriers in the isolation of co-occurring, closely related tree taxa, including at the early stages of speciation.

## Data availability

All data will be made available in Dryad upon acceptance of the manuscript for publication.

## Supporting Information

**Table S1** Site attributes of donor and maternal populations on Hawai’i Island.

**Table S2** Results from the best-fit linear mixed-effects model for the number of hand-pollinated flowers resulting in mature fruits in field crosses with 21 maternal trees of polymorpha.

**Table S3** AIC values used for model selection in the analyses of the number of weeks to maturity of hand-pollinated fruits on maternal trees of polymorpha on Hawaii Island.

**Table S4** Results from a linear mixed-effects model for the maximum number of germinants per fruit resulting from field crosses with 21 maternal trees of polymorpha.

**Table S5** AIC values used for model selection in the analyses of the maximum number of germinants per fruit and time to maximum germination from experimental crosses with 21 maternal trees of polymorpha on Hawaii Island.

**Table S6** AIC values used for model selection in the analyses of the maximum number of germinants per fruit and time to maximum germination from self- and open-pollinations on maternal trees of polymorpha on Hawaii Island.

**Fig. S1** Mean ± 1 SE fruit maturation time in weeks of inflorescences of each cross type at each of 21 maternal trees of polymorpha.

**Fig. S2** Variation in the maximum number of germinants per fruit resulting from controlled crosses on 21 maternal trees of polymorpha in the field, and self- and open-pollinated fruits from the same 21 trees.

**Fig. S3** Variation in the timing of seed germination for fruits resulting from controlled crosses on 21 maternal trees of polymorpha in the field, and self- and open-pollinated fruits from the same 21 trees.

**Fig. S4** Boxplots of variation in morphology (PC2 and PC3 scores) among 2-year-old seedings representing four genotypes produced through controlled crosses on 21 maternal trees of polymorpha.

**Fig. S5** The percentage of greenhouse-raised trees of four Hawaii-Island varieties of *M. polymorpha* and four F1 hybrids that reached the flowering stage each year over 12 years.

**Fig. S6** Boxplots for the percentage fruit set resulting from controlled outcrosses among greenhouse and common-garden trees of four Hawaii Island varieties of *M. polymorpha* and four of their F1 hybrids.

**Fig. S7** Boxplots for maturation time for fruits resulting from controlled outcrosses among greenhouse and common-garden trees of four Hawaii Island varieties of *M. polymorpha* and four of their F1 hybrids.

**Fig. S8** Percent normal pollen of four Hawaii-Island varieties of *M. polymorpha* and four F1 hybrids raised in the greenhouse/common garden.

**Fig. S9** Variation in proximal pollen-tube-bundle density in controlled outcrosses among greenhouse and common-garden trees of four Hawaii Island varieties of *M. polymorpha* and four of their F1 hybrids.

