## Supplementary Materials for "The importance of postzygotic barriers at the early stages of speciation in trees"

The following Supporting Material is available for this article:

**Fruit set and seed germination of self and open pollinations on polymorpha:** METHODS: To assess self-fertility, all 21 maternal trees were self-pollinated at ≥15 emasculated flowers, following the procedures used for outcrossed pollinations. Additional untreated and unbagged (i.e., open-pollinated) inflorescences on all maternal trees were also tagged as available at the start of anthesis. Recording of fruit set rates of the self- and open-pollinated inflorescences and seed germination rates of the self-pollinated inflorescences followed the methods for the outcrossed inflorescences. *Analysis:* The rate and timing of both fruit set and seed germination were analyzed using the analogous models for outcrossed inflorescences, but with polymorpha-outcrossed inflorescences from the same maternal tree included as the control.

RESULTS: *Fruit-set rate and maturation time—*The fruit set rate of the 564 self-pollinated flowers was 56.5%. Fruits derived from self-pollen were only 24% as likely to develop mature fruits as the control (z=-2.73 P=0.006). In contrast, fruit-set rate of the 537 open-pollinated flowers was high (81.3%) and not different from that of the control flowers (z=-0.56, P=0.57). Both self- and open-pollinated fruits matured over a mean of 46 weeks, which did not differ from the control.

*Rate and timing of seed germination—*The maximum number of germinants per fruit from self pollinations varied from 1 to 95. Germination of self-pollinated fruits was significantly lower (22.67 +/- 2.46 (SE) germinants/fruit) than germination of control fruits (104.83 +/- 10.09 germinants/fruit; t=-5.694; P<0.001). Seed germination of open-pollinated fruits (34.56 +/- 4.31) was also lower than that of the control fruits (t=-4.914; P<0.001), consistent with pollen limitation at the high-elevation edge of *Metrosideros*’ range. The number of weeks to peak seed germination was lower for self- and open-pollinated fruits (2.94 +/- 0.37 (SE) weeks; t=-3.077; P=0.003; and 3.07 +/- 0.34 weeks; t=-2.740; P=0.008, respectively) compared to the control (3.77 +/- 0.42; Table S6).

**Table S1** Site attributes of donor and maternal populations on Hawai’i Island.  Eight donor (♂) populations (two sites per variety: *M. polymorpha* var. *glaberrima* (G), *M. polymorpha* var. *newellii* (N), *M. polymorpha* var. *incana* (I) and *M. polymorpha* var. *polymorpha* (P)) were used in an experimental cross with a single maternal population (♀P).

| Population | Site Name | Latitude | Longitude | Altitude (m) |
| --- | --- | --- | --- | --- |
| ♂ G1 | Stainback Highway | 19.571 | 155.205 | 966 |
| ♂ G2 | Upper Wailuku River | 19.709 | -155.270 | 1140 |
| ♂ I1 | Kapoho Kalapana Rd. | 19.479 | -154.833 | 21 |
| ♂ I2 | Mohouli St. | 19.703 | -155.096 | 194 |
| ♂ N1 | Wailuku River | 19.708 | -155.234 | 907 |
| ♂ N2 | Kalohewahewa River | 19.715 | -155.186 | 619 |
| ♂ P1 | Lower Observatory Rd. | 19.675 | 155.467 | 2019 |
| ♂ P2 | Mauna Loa Cabin Trail | 19.494 | 155.386 | 2066 |
| ♀ P | Upper Observatory Rd. | 19.607 | -155.468 | 2374 |

**Table S2** Results from the best-fit linear mixed-effects model for the number of hand-pollinated flowers resulting in mature fruits in field crosses with 21 maternal trees of polymorpha; control = polymorpha.

|  | Fruit set | | | | |
| --- | --- | --- | --- | --- | --- |
| Pollen donor | Mean | SE | Log-odds ratio | *x*-value | P-value |
| control | 0.79 | 0.04 | 0.84 | 4.43 | - |
| glaberrima | 0.74 | 0.03 | 0.78 | -1.54 | 0.125 |
| incana | 0.75 | 0.05 | 0.82 | -0.34 | 0.719 |
| newellii | 0.71 | 0.04 | 0.71 | -2.74 | 0.006 |

**Table S3** AIC values used for model selection in the analyses of the number of weeks to maturity of hand-pollinated fruits on maternal trees of polymorpha on Hawaii Island.  Model 1 includes a random slope for each variety within individual maternal trees (random effect) with correlated intercepts.  In model 2 the intercepts vary among variety and among maternal trees (random effect) within variety (nested random effects), while model 3 allows each population a random intercept.

|  | AIC Values | |
| --- | --- | --- |
| Model | Weeks to maturity | Fruit set |
| null model | 232.0916 | 674.1603 |
| model 1 | 221.9163* | 211.3693* |
| model 2 | 226.5108 | 382.9759 |
| model 3 | 227.4062 | 501.5881 |

**Table S4** Results from a linear mixed-effects model for the maximum number of germinants per fruit (square-root-transformed for analysis) resulting from field crosses with 21 maternal trees of polymorpha; control = polymorpha.

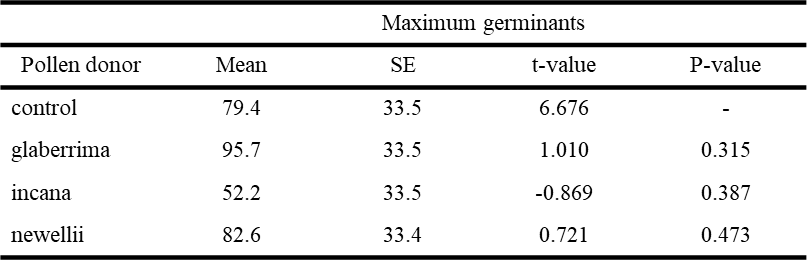

**Table S5** AIC values used for model selection in the analyses of the maximum number of germinants per fruit and time to maximum germination from experimental crosses with 21 maternal trees of polymorpha on Hawaii Island. Model 1 includes a random slope for each variety within individual maternal trees (random effect) with correlated intercepts. In model 2 the intercepts vary among variety and among maternal trees (random effect) within variety (nested random effects), while model 3 allows each population a random intercept.

|  | AIC Values | |
| --- | --- | --- |
| Model | Seed germination | Time to maximum germinants |
| null model | 2193.574 | 1725.114 |
| model 1 | 2176.814 | 1644.555 |
| model 2 | 2063.321* | 1586.842* |
| model 3 | 2222.993 | 1651.743 |

**Table S6** AIC values used for model selection in the analyses of the maximum number of germinants per fruit and time to maximum germination from self- and open-pollinations on maternal trees of polymorpha on Hawaii Island.  Model 1 includes a random slope for each variety within individual maternal trees (random effect) with correlated intercepts.  In model 2 the intercepts vary among variety and among maternal trees (random effect) within variety (nested random effects), while model 3 allows each population a random intercept.

|  | AIC Values | |
| --- | --- | --- |
| Model | Seed germination | Time to maximum germinants |
| null model | 1393.438 | 971.864 |
| model 1 | 1336.547 | 951.843 |
| model 2 | 1215.607* | 857.864* |
| model 3 | 1315.002 | 972.059 |

**Fig. S1** Mean ± 1 SE fruit maturation time in weeks of inflorescences of each cross type at each of 21 maternal trees of polymorpha.  Pollen-donor types: G=glaberrima, I=incana, N=newellii, P=polymorpha.  Fruit maturation time varied among maternal trees, but the pattern in maturation time among cross types was not consistent across maternal trees.

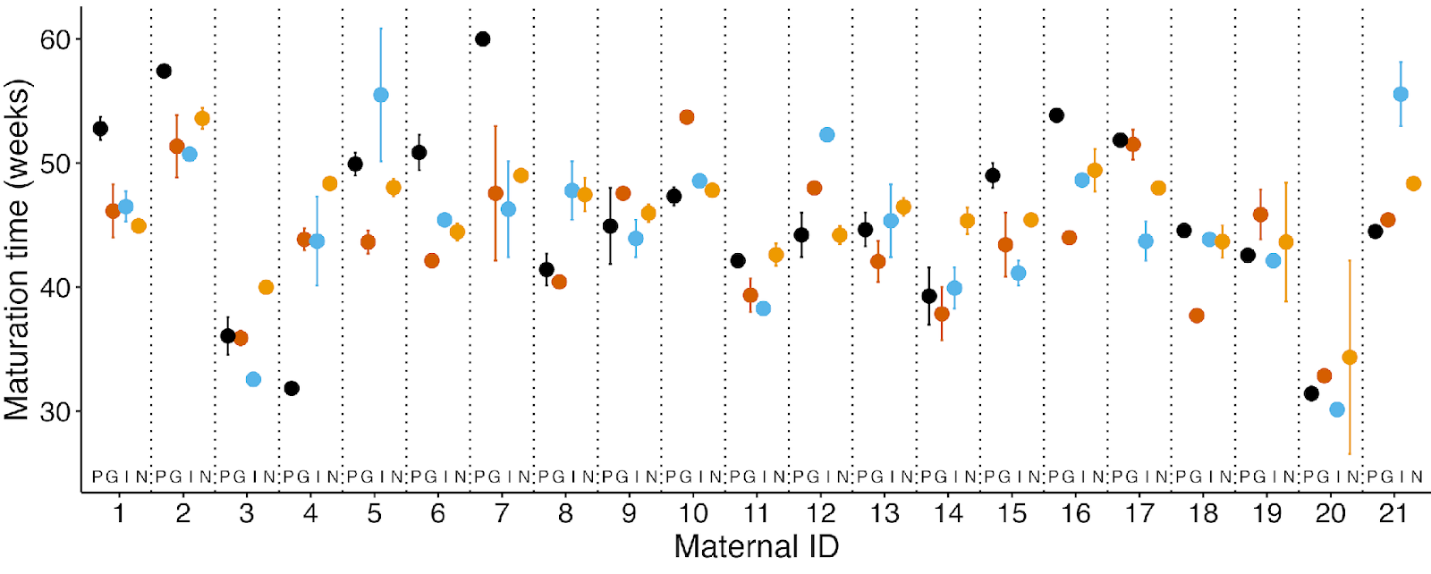

**Fig. S2** Variation in the maximum number of germinants per fruit resulting from (left) controlled crosses on 21 maternal trees of polymorpha in the field, and (right) self- and open-pollinated fruits from the same 21 trees.  Shown are means (filled circles), medians (lines), and individual data points (open circles).  Sample sizes are along the top; 10 maternal trees received pollen from both N populations.  Pollen-donor variety codes are as in Fig. S1; control=polymorpha.  * indicates significant difference from the control.

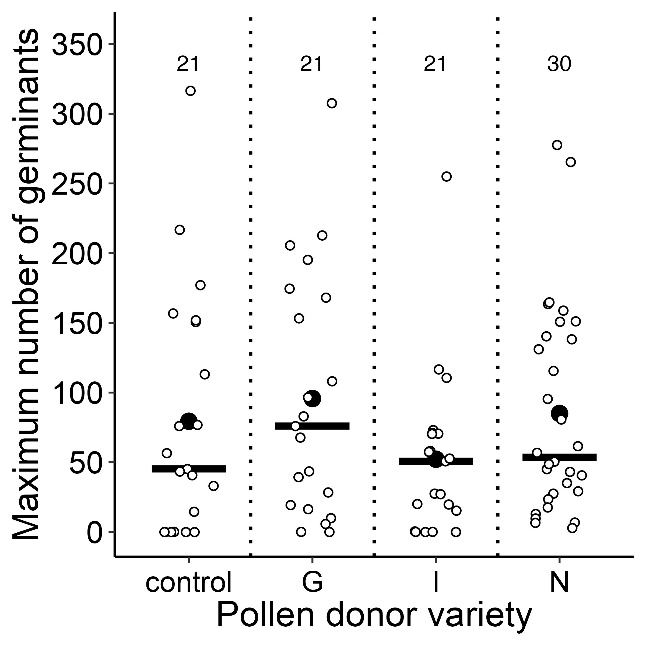

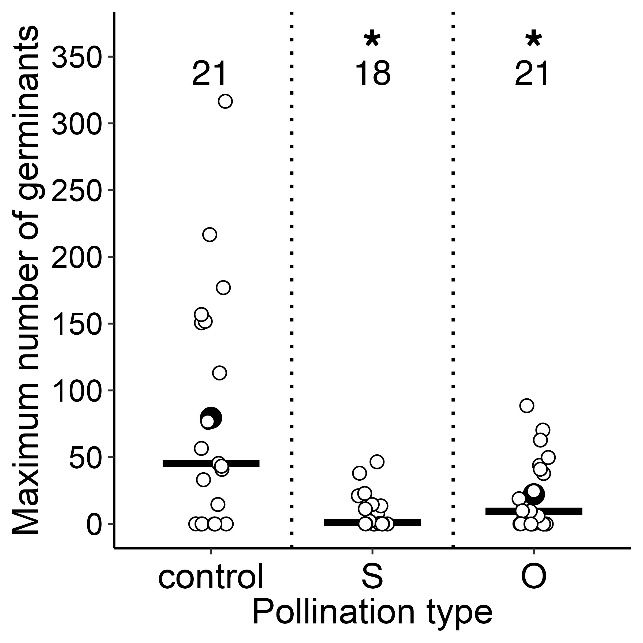

**Fig. S3** Variation in the timing of seed germination (weeks to maximum germinants) for fruits resulting from (left) controlled crosses on 21 maternal trees of polymorpha in the field, and (right) self- and open-pollinated fruits from the same 21 trees.  Shown are means (filled circles), medians (lines), and individual data points (open circles).  Sample sizes are along the top; 10 maternal trees received pollen from both N populations.  Pollen-donor variety codes are as in Fig. S1; control=polymorpha.  * indicates significant difference from the control.

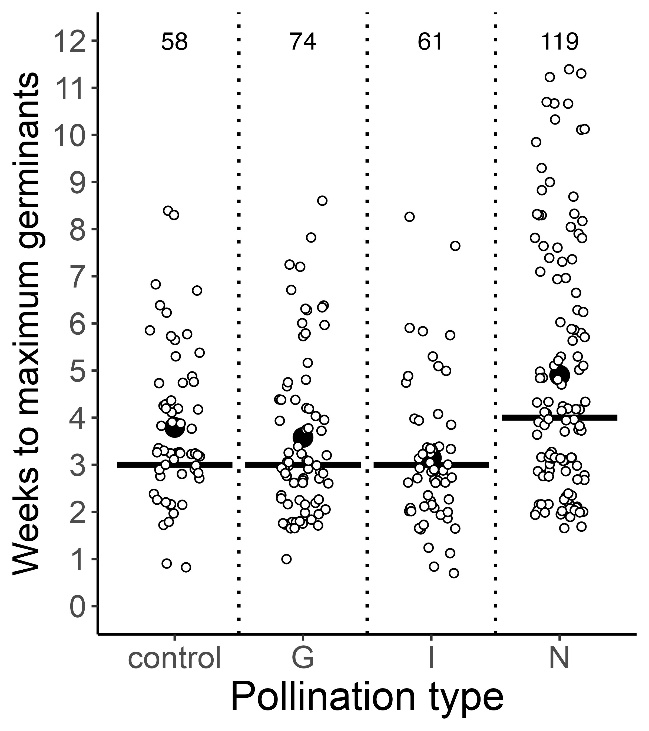

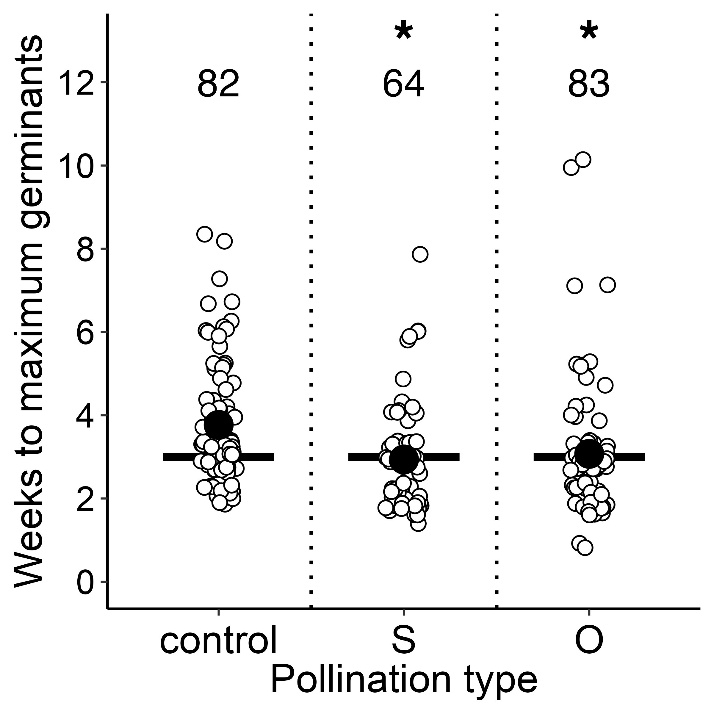

**Fig. S4** Boxplots of variation in morphology (PC2 and PC3 scores) among 2-year-old seedings representing four genotypes produced through controlled crosses on 21 maternal trees of polymorpha. Genotypes: control=polymorpha, GP=glaberrimaXpolymorpha, IP=incanaXpolymorpha, NP=newelliiXpolymorpha.  Circles indicate means; lines indicate medians.  * indicates significant difference from the control. Mean scores for PC2 (19.1%), which largely represent a trade-off between leaf and petiole lengths versus numbers of leaves and stems, were greater for all three F_1_ genotypes relative to the controls (t=7.53-11.19, P<0.001), reflecting the lower number (except for NP) of larger leaves of the F_1_ hybrids (Fig. 6). Mean scores of PC3 (14.4%), largely representing pubescence, differed significantly between the controls and both the GP and NP F_1_s (t=-2.61, P=0.009; t=-4.19, P<0.001, respectively; Fig. 6), consistent with the glabrousness of glaberrima and newellii (Fig. 2).

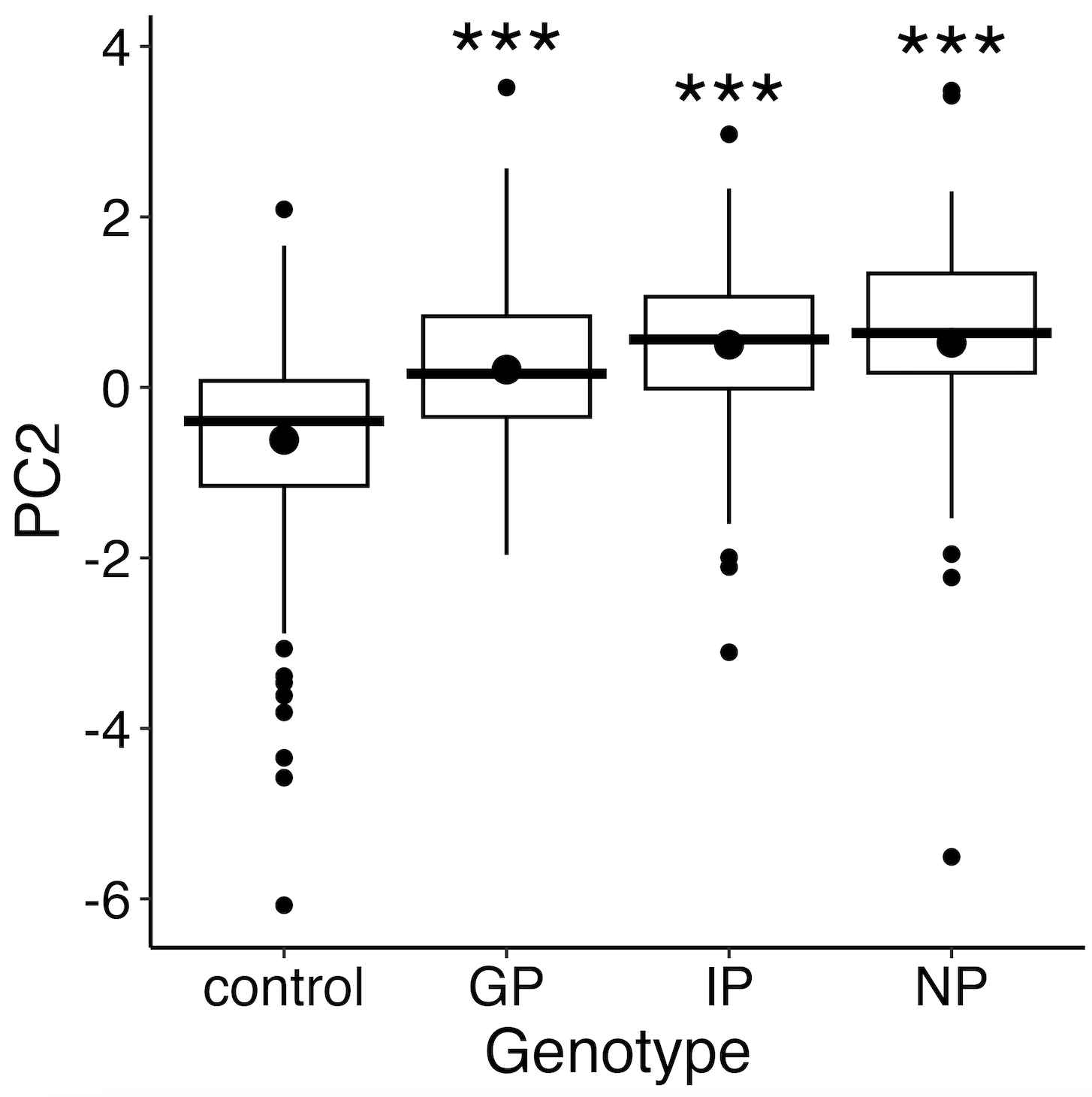

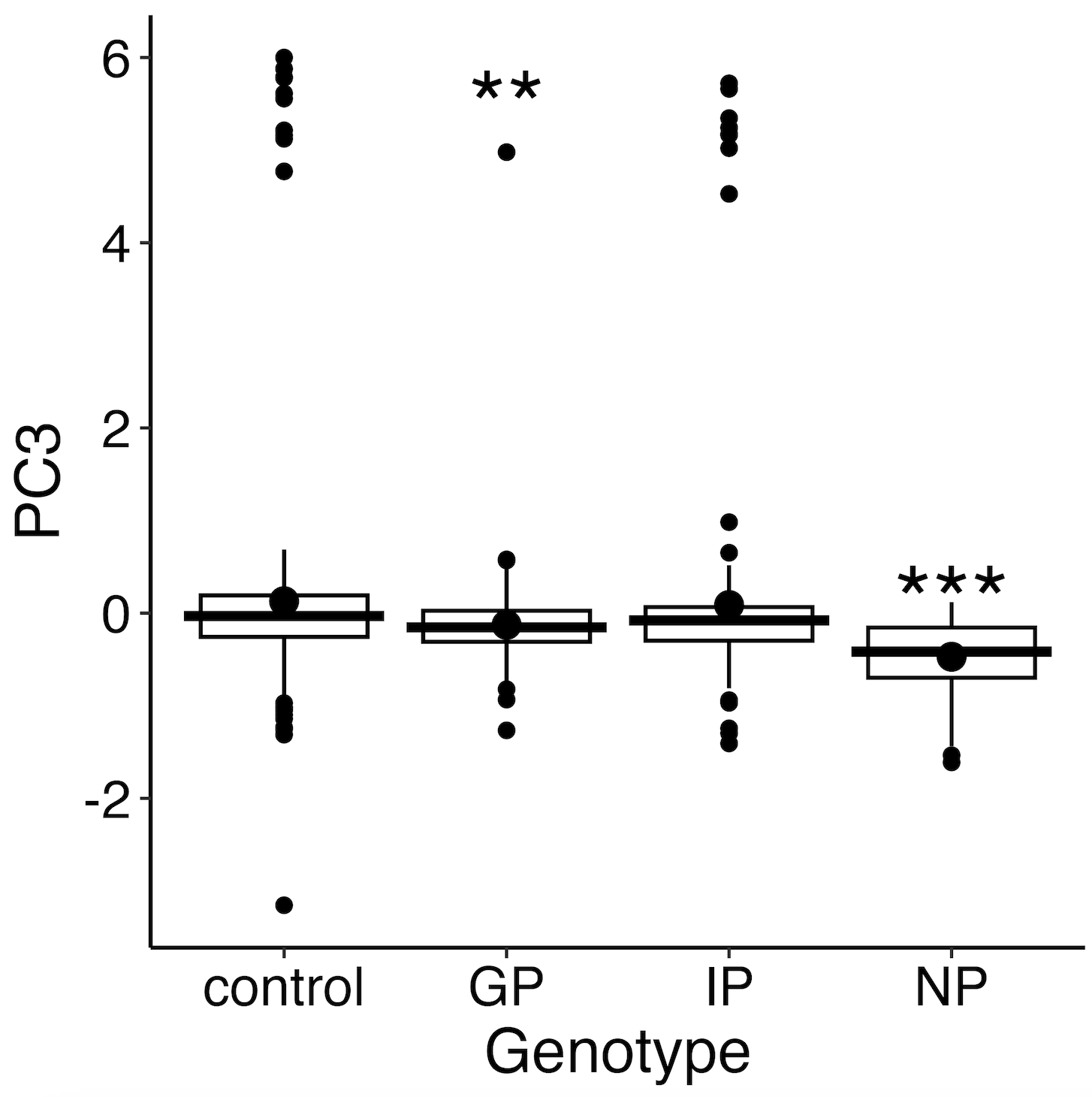

**Fig. S5** The percentage of greenhouse-raised trees of four Hawaii-Island varieties of *M. polymorpha* (G, I, N and P; see Fig. S1) and four F_1_ hybrids (GI, GP, IP, and NP) that reached the flowering stage each year (from the seed stage) over 12 years (i.e., maturation rate).

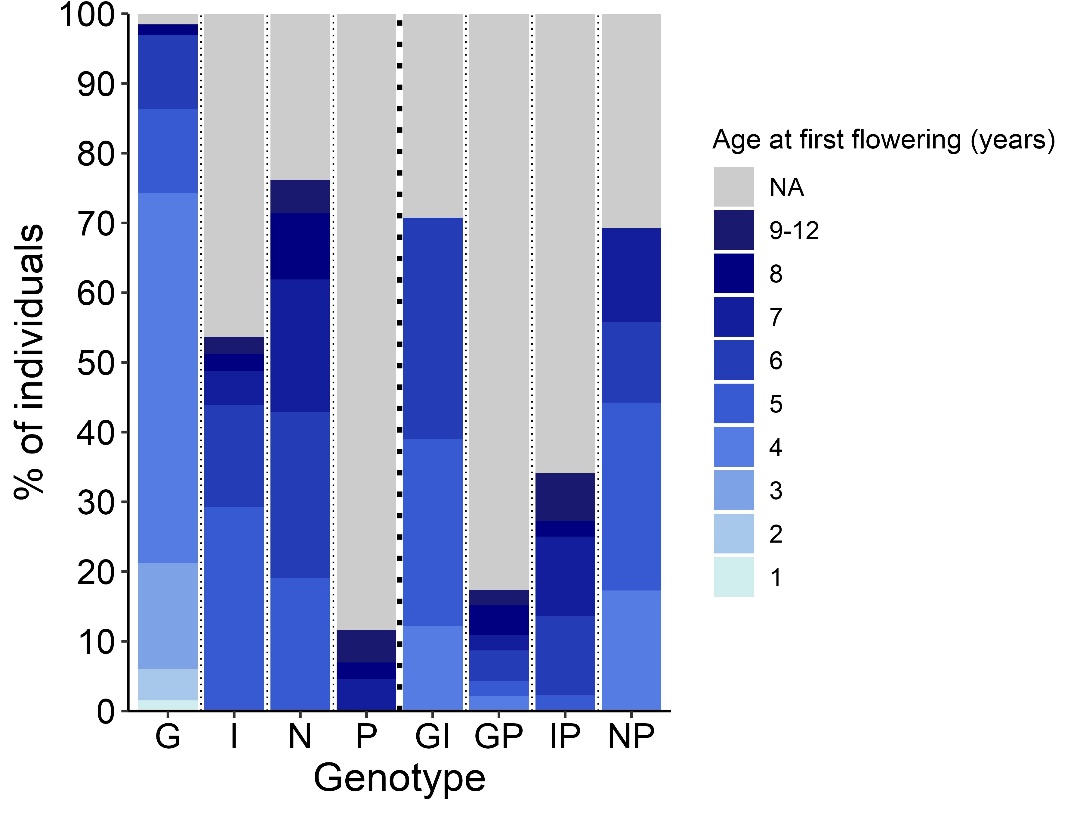

**Fig. S6** Boxplots for the percentage fruit set resulting from controlled outcrosses among greenhouse and common-garden trees of four Hawaii Island varieties of *M. polymorpha* (G, I, N and P; see Fig. S1) and four of their F_1_ hybrids (GI, GP, IP, and NP).  *Left*: crosses are pooled (among pollen-donor genotypes) within maternal genotypes; *right*: crosses are pooled (among maternal-tree genotypes) within pollen-donor genotypes.

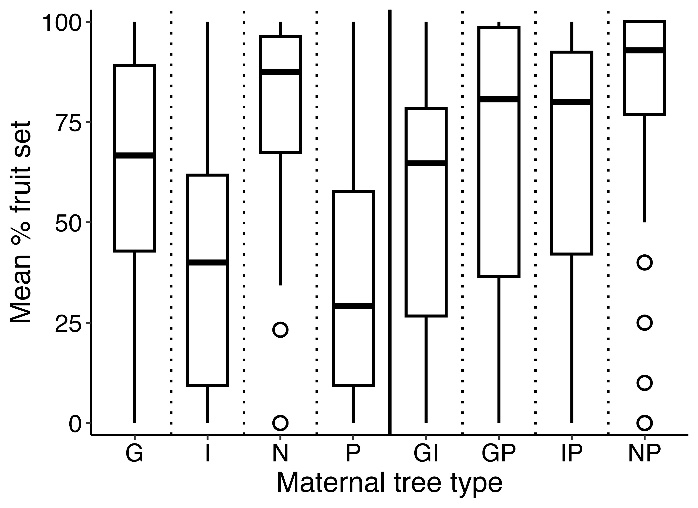

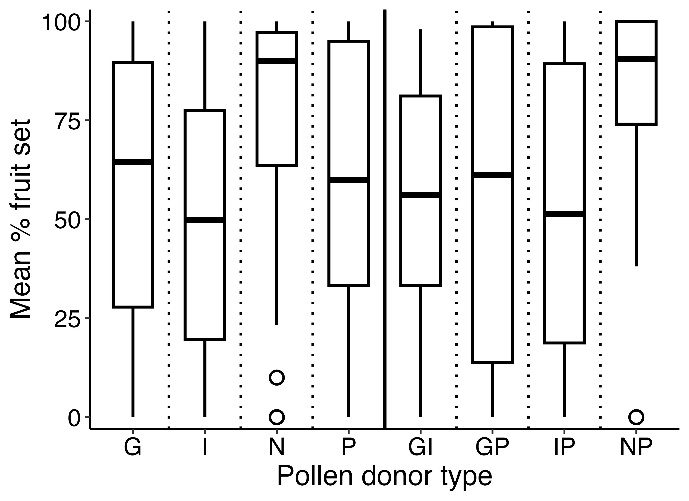

**Fig. S7** Boxplots for maturation time (in weeks) for fruits resulting from controlled outcrosses among greenhouse and common-garden trees of four Hawaii Island varieties of *M. polymorpha* (G, I, N and P; see Fig. S1) and four of their F_1_ hybrids (GI, GP, IP, and NP).  *Left*: crosses are pooled (among pollen-donor genotypes) within maternal genotypes; *right*: crosses are pooled (among maternal-tree genotypes) within pollen-donor genotypes.

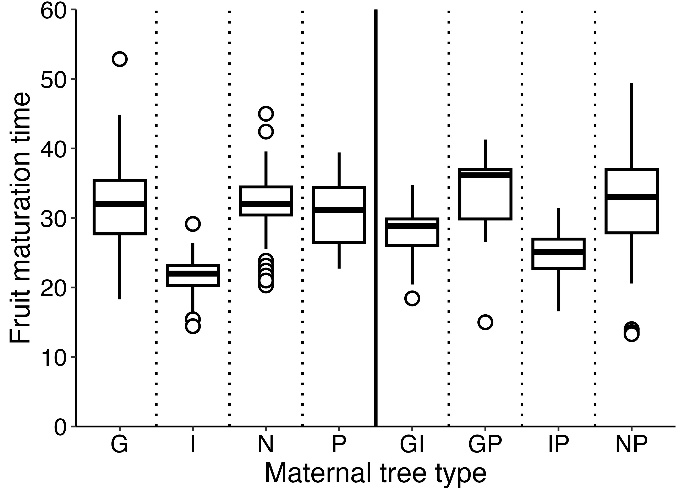

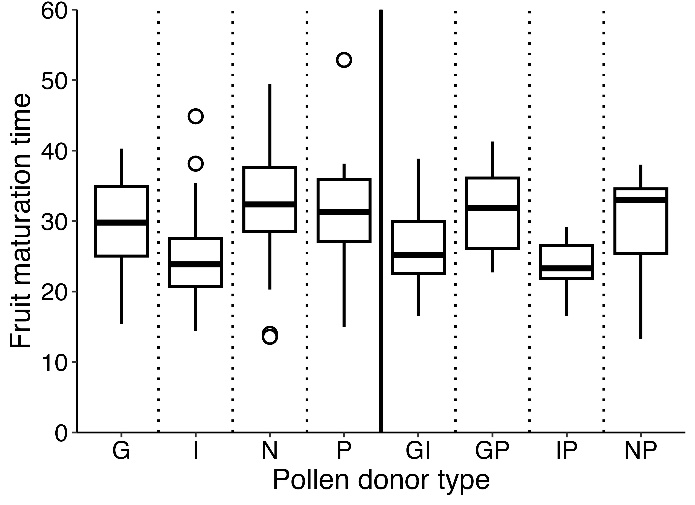

**Fig. S8** Percent normal pollen (stained with lactophenol cotton blue) of four Hawaii-Island varieties of *M. polymorpha* (G, I, N and P; see Fig. S1) and four F_1_ hybrids (GI, GP, IP, and NP) raised in the greenhouse/common garden.  Shown are medians (lines) and individual data points (open circles).  Sample sizes are along the top.

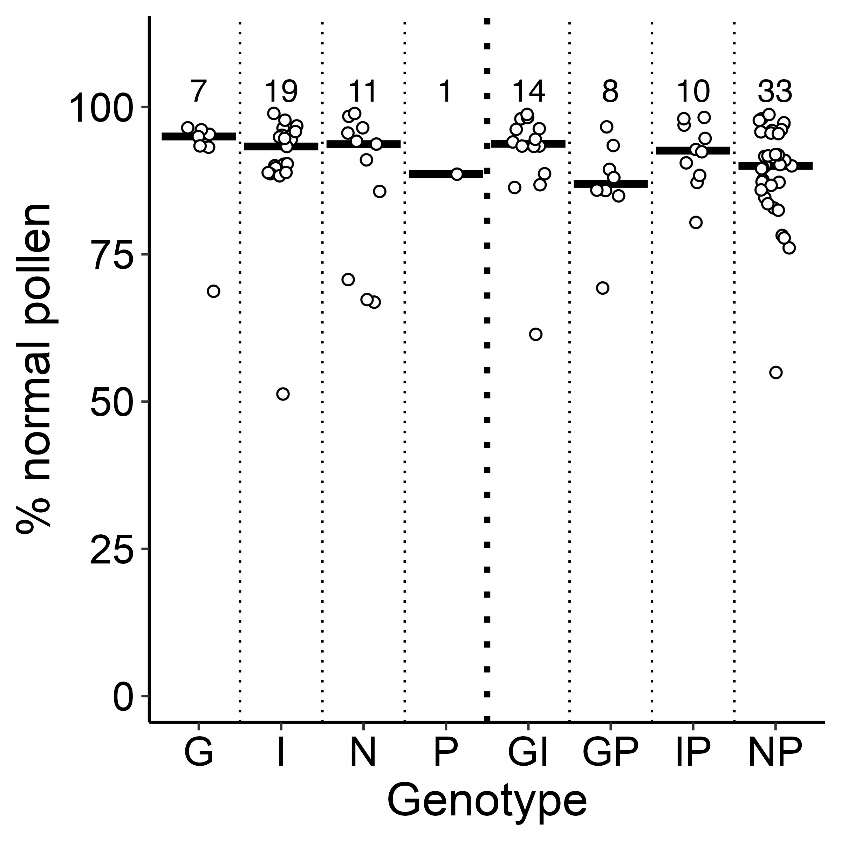

**Fig. S9** Variation in proximal pollen-tube-bundle (PTB) density in controlled outcrosses among greenhouse and common-garden trees of four Hawaii Island varieties of *M. polymorpha* (G, I, N and P; see Fig. S1) and four of their F_1_ hybrids (GI, GP, IP, and NP).  *Left*: crosses are pooled (among pollen-donor genotypes) within maternal genotypes; *right*: crosses are pooled (among maternal-tree genotypes) within pollen-donor genotypes.  Sample sizes are along the top.  Typically, five styles per cross were examined 7 days post-pollination.  Proximal PTB density was strongly representative of PTB density throughout the full length of the style.

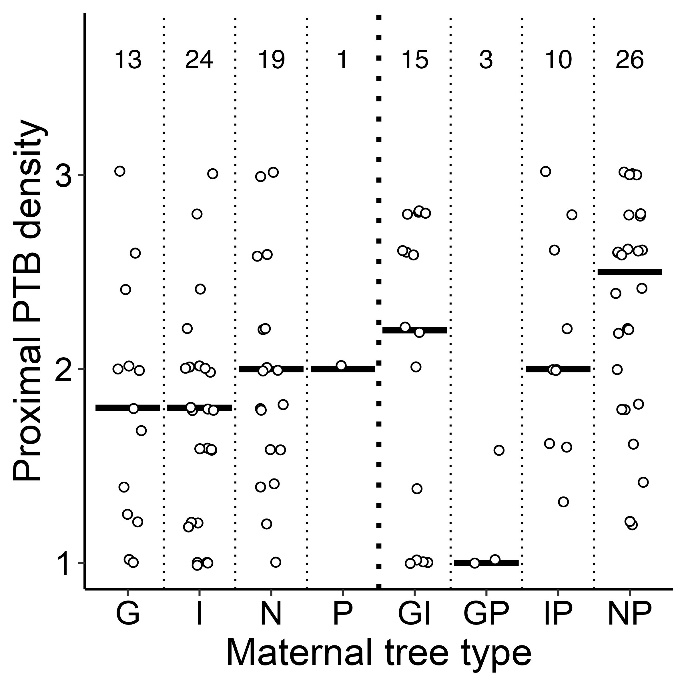

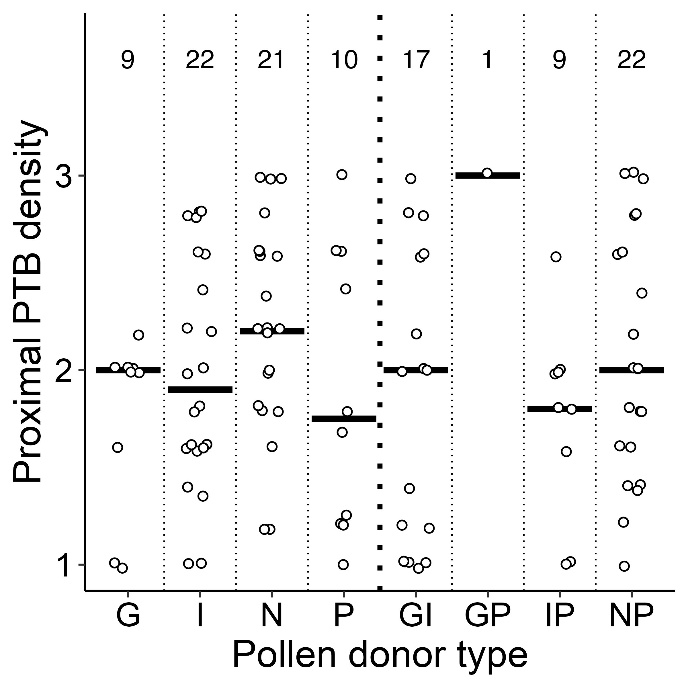
